# Disease related mutations in PI3Kγ disrupt regulatory C-terminal dynamics and reveals a path to selective inhibitors

**DOI:** 10.1101/2020.12.01.406264

**Authors:** Manoj K Rathinaswamy, Zied Gaieb, Kaelin D Fleming, Chiara Borsari, Noah J Harris, Brandon E Moeller, Matthias P Wymann, Rommie E Amaro, John E Burke

## Abstract

Class I Phosphoinositide 3-kinases (PI3Ks) are master regulators of cellular functions, with the p110*γ* subunit playing a key role in immune signalling. PI3K*γ* is a key factor in inflammatory diseases, and has been identified as a therapeutic target for cancers due to its immunomodulatory role. Using a combined biochemical/biophysical approach, we have revealed insight into regulation of kinase activity, specifically defining how immunodeficiency and oncogenic mutations of R1021 in the c-terminus can inactivate or activate enzyme activity. Screening of small molecule inhibitors using HDX-MS revealed that activation loop binding inhibitors induce allosteric conformational changes that mimic those seen for the R1021C mutant. Structural analysis of clinically advanced PI3K inhibitors revealed novel binding pockets that can be exploited for further therapeutic development. Overall this work provides unique insight into the regulatory mechanisms that control PI3K*γ* kinase activity, and shows a framework for the design of PI3K isoform and mutant selective inhibitors.

## Introduction

The phosphoinositide 3-kinase (PI3K) family of enzymes are central regulators of growth, proliferation, migration, and metabolism in a plethora of cells and tissues [1, 2]. PI3Ks are lipid kinases that generate the lipid second messenger phosphatidylinositol 3,4,5 trisphosphate (PIP_3_), which is utilised downstream of cell surface receptors to regulate growth, metabolism, survival, and differentiation [1]. PIP_3_, is generated by four distinct class I PI3K catalytic isoforms separated into two groups (class IA [p110*α*, p110*β*, p110*δ*], and class IB [p110*γ*] (sometimes referred to as PI3K*α*, PI3K*β*, PI3K*δ*, and PI3K*γ* catalytic subunit)). The primary difference between class IA and class IB PI3Ks is their association with specific regulatory subunits, with class IA binding p85-like regulatory subunits encoded by *PIK3R1, PIK3R2, PIK3R3*, and PI3K*γ* forming complexes with either a p101 or p84 (also called p87^PIKAP^) adaptor subunit [3–5]. The four isoforms of class I PI3K have distinct expression profiles, with PI3K*α* and PI3K*β* being ubiquitously expressed, and PI3K*δ* and PI3K*γ* being mainly localised in immune cells [1]. All PI3K isoforms have been implicated in a variety of human diseases, including cancer, immunodeficiencies, inflammation, and developmental disorders [6–8].

The class IB PI3K*γ* isoform encoded by *PIK3CG* is a master regulator of immune cell function. It plays important roles in the regulation of myeloid (macrophages, mast cells, neutrophils) and lymphoid (T cells, B cells, and Natural Killer cells) derived immune cells [9–11]. PI3K*γ* regulates immune cell chemotaxis [11–13], cytokine release [14, 15], and generation of reactive oxygen species[11], which are important processes in both the innate and adaptive immune systems. The ability of PI3K*γ* to mediate multiple immune cell functions is controlled by its activation downstream of numerous cell surface receptors, including G-protein coupled receptors (GPCRs)[16], the IgE/Antigen receptor[14], receptor tyrosine kinases (RTKs) [17], and the Toll-like receptors (TLRs) [18, 19]. Activation of PI3K*γ* downstream of these stimuli are potentiated by their p84 and p101 regulatory subunits [5,18,20–22]. This is distinct from the roles of regulatory subunits in class IA PI3Ks, which act as potent inhibitors of p110 catalytic activity[23]. In mouse models, loss of PI3K*γ* either genetically or pharmacologically is protective in multiple inflammatory diseases including cardiovascular disease [10], arthritis [9], Lupus [24], asthma [15], pulmonary inflammation and fibrosis [25, 26], and metabolic syndrome [27]. PI3K*γ* is also a driver of pancreatic ductal adenocarcinoma progression through immunomodulatory effects [28], and targeting PI3K*γ* in the immune system in combination with checkpoint inhibitors has shown promise as an anti-cancer therapeutic [29, 30].

Extensive biophysical and biochemical assays have identified many of the molecular mechanisms underlying PI3K*γ* regulation. The enzyme is composed of five domains, a putative uncharacterized adaptor binding domain (ABD), a Ras binding domain (RBD), a C2 domain, a helical domain, and a bi-lobal lipid kinase domain [31] (Fig. 1A). PI3K*γ* activation is primarily mediated by G*βγ* subunits downstream of GPCR signalling, through a direct interaction of G*βγ* with the C2-helical linker of PI3K*γ* [21]. Activation of PI3K*γ* by G*βγ* requires a secondary interaction between G*βγ* and regulatory subunits for physiologically relevant activation [4], with the free p110*γ* subunit in cells having no detectable activation downstream of GPCR activation [32]. In addition, PI3K*γ* activation can be facilitated by Ras GTPases interacting with the RBD [33], with the same interface putatively also mediating activation by Rab8 [19]. Experiments exploring a novel type II-like kinase inhibitor that targets an active conformation of PI3K*γ* revealed novel molecular aspects of regulation involving the C-terminal regulatory motif of the kinase domain, which is composed of the k*α*7, 8, 9, 10, 11, 12 helices that surround the activation loop, and keep the enzyme in an inhibited state [34] (Fig. 1B). The k*α*10, k*α*11, and k*α*12 helices are sometimes referred to as the regulatory arch [35]. Inhibition mediated by the C-terminal regulatory motif is conserved through all class I PI3Ks, although for all other isoforms, this inhibited conformation requires interactions with a p85 regulatory subunit (Fig. S1) [8]. In PI3K*γ* this inhibited conformation is proposed to be maintained by a Tryptophan lock, where W1080 maintains a closed conformation of the membrane binding C-terminal k*α*12 helix, leading to an inactive conformation of the activation loop [34] (Fig. 1B+C).

**Fig 1.**
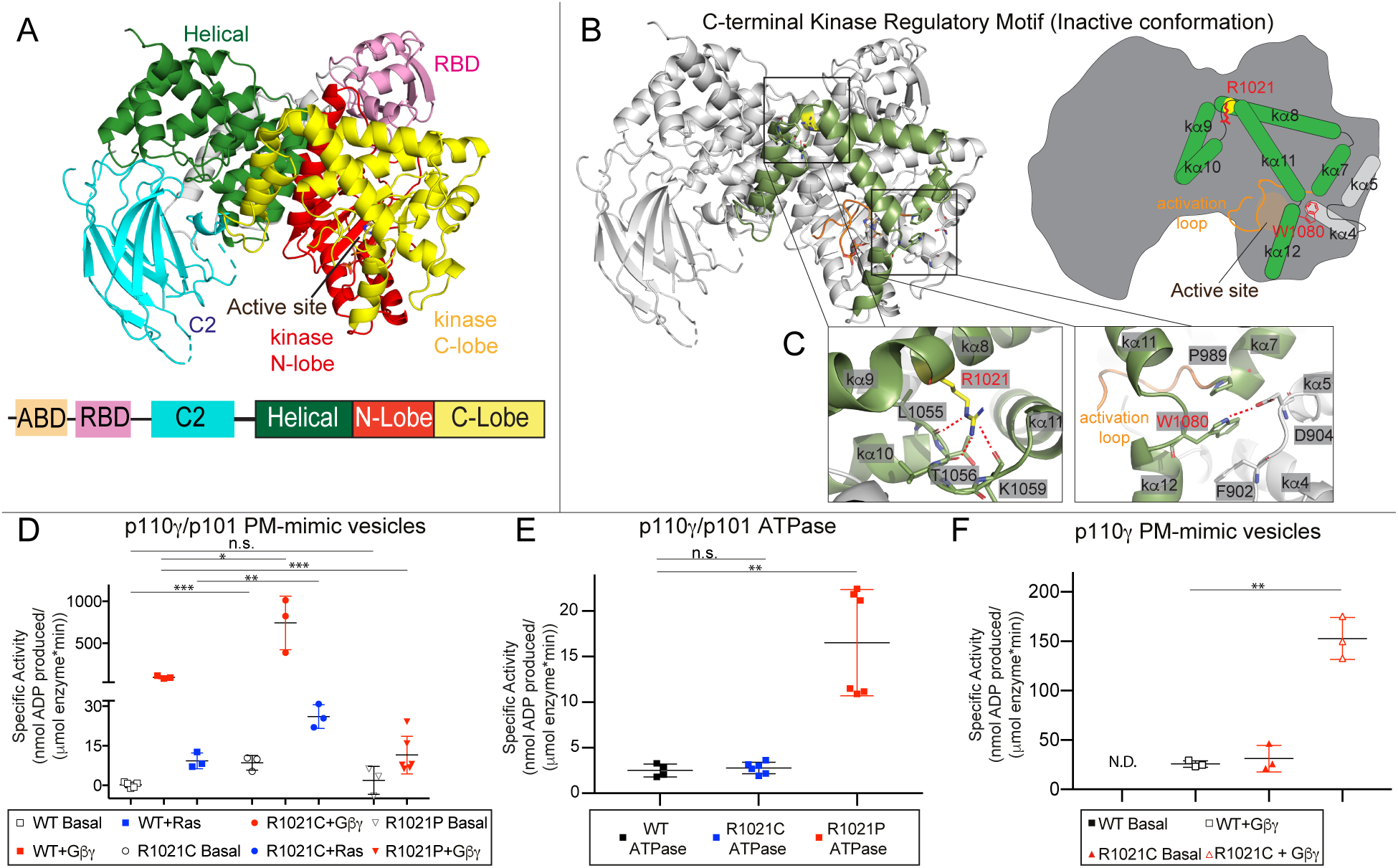
Mutations in the regulatory C-terminal motif of the kinase domain alter PI3Kγ activity. **A.** Domain architecture of p110*γ* (PDB ID: 6AUD) [54], with the domain schematic shown beneath. **B.** Model of the C-terminal regulatory motif of the kinase domain of p110*γ*. The helices that make up the regulatory arch (k*α*10, 11,12) and those that pack against them (k*α*7, 8, 9) are highlighted in green both in the structural model and cartoon schematic. **C.** A close up of the W1080 ‘Tryptophan lock’ interaction with k*α*7 and the k*α*4-k*α*5 loop which maintains an inhibited conformation is shown, as well as the interaction of the R1021 residue with residues on the k*α*10-k*α*11 loop. **D.** Lipid kinase activity assays testing the activity of WT, R1021C, and R1021P p110*γ*/p101 WT basally and in the presence of lipidated G*βγ* and HRas. Experiments were carried out with 50-3000 nM kinase, 1500 nM Ras, 1500 nM G*βγ*, all in the presence of 100 µM ATP and 1 mg/mL PM-mimic vesicles [5% phosphatidylinositol 4,5 bisphosphate (PIP_2_), 20% phosphatidylserine (PS), 10% phosphatidyl choline (PC), 50% phosphatidylethanolamine (PE), 10% Cholesterol, 5% sphingomyelin (SM)). **E.** Activity assays testing the intrinsic ATPase activity (ATP conversion in the absence of membrane substrate) for wild type and mutant p110*γ*/p101 complexes. **F.** Lipid kinase activity assays testing the activity of WT and R1021C for the free p110*γ* catalytic subunit with and without lipidated G*βγ*. Lipid kinase activity was generated by subtracting away non-specific ATPase activity, for unstimulated WT p110*γ* there was no detectable lipid kinase activity above basal ATPase activity (N.D.). For panels D-F, every replicate is plotted, with error shown as standard deviation (n=3-6). Two tailed p-values represented by the symbols as follows: ***<0.001; **<0.01; *<0.05; N.S.>0.05.

Disruption of PI3K signalling by either activating or inactivating mutations and deletions are involved in multiple human diseases. Overexpression of any activated class I PI3K isoform can lead to oncogenic transformation [36], although PI3K*α* is the most frequently mutated in human disease. Activating PI3K*α* mutations are linked to both cancer [37, 38] and overgrowth disorders [39], with activating PI3K*δ* mutations linked to primary immuno-deficiencies [40–42]. A high proportion of these activating mutations cluster to the C-terminal regulatory motif of PI3Ks. Multiple PI3K*γ* mutations have been identified in cancer patients [43–45], although at a lower frequency than PI3K*α* mutations. It would be expected that these mutations are activating, although this has not been fully explored. Intriguingly, PI3K*γ* loss of function mutations in the C-terminal regulatory motif (R1021P, N1085S) have been identified in patients with immunodeficiencies [46, 47] (Fig. 1B+C). PI3K mediated diseases being caused by both activating and inactivating mutations, highlights the critical role of maintaining appropriate PIP_3_ levels for human health.

The involvement of activated PI3K signalling in multiple diseases has motivated class I PI3K inhibitor development. There is, however, toxicity effects associated with compounds that target all PI3K isoforms by mechanism-based adverse side effects [48], driving the development of isoform selective inhibitors. These efforts have led to multiple clinically approved inhibitors of PI3K*α* and PI3K*δ* [49–51]. The critical role of PI3K*γ* in inflammation and the tumour microenvironment has stimulated development of PI3K*γ* specific inhibitors. Two main strategies for generating PI3K*γ* selective ATP-competitive inhibitors have been established: i) targeting PI3K*γ* specific regions outside of the ATP binding pocket to reach regions not conserved among PI3K isoforms [52, 53], and ii) targeting selective PI3K*γ* conformational changes [34]. Intriguingly, the conformational selective PI3K*γ* inhibitors appear to target its putatively activated conformation.

The parallel discovery of disease linked mutations in the C-terminal regulatory motif, and conformational selective PI3K*γ* inhibitors that cause altered dynamics of the C-terminus led us to investigate the underlying molecular mechanisms. Using a combined biochemical and biophysical approach, we characterized the dynamic conformational changes caused by the loss of function R1021P mutation, as well as a putative oncogenic R1021C mutation identified in Catalogue of Somatic Mutations in Cancer database [COSMIC [45]]. A screen of a number of PI3K*γ* selective and pan-PI3K inhibitors revealed that many of these molecules induced allosteric conformational changes in PI3K*γ*. A combined X-ray crystallography and hydrogen deuterium exchange mass spectrometry (HDX-MS) approach showed that inhibitor interactions with the activation loop mediates allosteric conformational changes. Intriguingly, similar conformational changes occurred for both the R1021C mutant and upon binding certain inhibitors, with lipid kinase assays revealing an increased potency of these inhibitors towards the activated PI3K mutant. Overall, this work provides a unique insight into how mutations alter PI3K*γ* regulation, and pave the way to novel strategies for isoform and mutant selective PI3K inhibitors.

## Results

### R1021P and R1021C mutations alter the activity of PI3Kγ

The recent discovery of an inactivating disease-linked mutation in *PIK3CG* located near the C-terminus of the kinase domain (R1021P) in immunocompromised patients led us to investigate the molecular mechanism of this mutation. Intriguingly, this same residue is found to be mutated in the COSMIC database (R1021C) [45]. To define the effect of these mutations on protein conformation and biochemical activity, we generated them recombinantly in complex with the p101 regulatory subunit. Both the p110*γ* R1021C and R1021P complexes with p101 eluted from gel filtration similar to wild-type p110*γ*, suggesting they were properly folded (Fig. S2). However, the yield of the R1021P complex with p101 was dramatically decreased relative to both wild-type and R1021C p110*γ*, indicating that this mutation may decrease protein stability, consistent with decreased p110*γ* and p101 expression in patient tissues [46]. We also generated the free R1021C p110*γ* subunit, however we could not express free p110*γ* R1021P, further highlighting that this mutation likely leads to decreased protein stability.

The R1021 residue forms hydrogen bonds with the carbonyl oxygens of L1055, T1056, and K1059 located in or adjacent to the regulatory arch helices k*α*10 and k*α*11 of PI3K*γ* (Fig 1C). Both R1021C and R1021P would be expected to disrupt these interactions, with the R1021P also expected to distort the secondary structure of the k*α*8 helix. The R1021P has been previously found to lead to greatly decreased lipid kinase activity *in vitro* [46]. To characterize these mutations, we carried out biochemical assays of wild-type, R1021C, and R1021P p110*γ*/p101 complexes against plasma membrane-mimic lipid vesicles containing 5% PIP_2_. Assays were carried out in the presence and absence of lipidated Gβ*γ* subunits, a potent p110*γ*/p101 activator. These assays revealed that p110*γ*/p101 R1021C was ∼8-fold more active than wild-type both basally and in the presence of Gβ*γ* (Fig. 1D). The R1021P complex showed greatly decreased Gβ*γ* stimulation compared to wild-type. Intriguingly, R1021P showed higher basal ATPase activity (non-productive turnover of ATP) compared to WT, revealing that it still has catalytic activity, but greatly decreased activity on lipid substrate (Fig. 1E). The R1021C mutant also showed a ∼8-fold increase in lipid kinase activity compared to wild-type when assaying the free 110*γ* subunit (Fig. 1F).

### R1021P and R1021C cause allosteric conformational changes throughout the regulatory C-terminal motif

We carried out hydrogen deuterium exchange mass spectrometry (HDX-MS) experiments to define the molecular basis for why two different mutations at the same site have opposing effects on lipid kinase activity. HDX-MS is a technique that measures the exchange rate of amide hydrogens, and as the rate is dependent on the presence and stability of secondary structure, it is an excellent probe of protein conformational dynamics [55]. HDX-MS experiments were performed on complexes of wild-type p110*γ*/p101, R1021C p110*γ*/p101, and R1021P p110*γ*/p101, as well as the free wild-type and R1021C p110*γ*. The coverage map of the p110*γ* and p101 proteins was composed of 153 peptides spanning ∼93% percent of the exchangeable amides (Table S1).

The R1021C and R1021P mutations led to significant changes in the conformational dynamics of the p110*γ* catalytic and p101 regulatory subunits (Fig. 2A-G). The R1021C mutation resulted in increased H/D exchange in the C2, helical and kinase domains of p110*γ*. Intriguingly, many of the changes in dynamics of the helical and kinase domains are similar to those observed upon membrane binding [21]. The largest differences occurred in the helices in the C-terminal regulatory motif (k*α*7-12) (Fig. 2C). A peptide spanning the C-terminal end of the activation loop and k*α*7 (976-992) showed increased exchange, with these changes primarily occurring at later timepoints of exchange (3000 s) (Fig. 2G). This is indicative of these regions maintaining secondary structure, although with increased flexibility. These increases in exchange for the R1021C mutant were conserved for the free p110*γ* subunit, although with larger increases in exchange compared to the p110*γ*/p101 complex (Fig. S3). Previous HDX-MS analysis of the regulatory mechanisms of class IA PI3Ks has revealed that increased dynamics of the activation loop occurs concurrently with increased lipid kinase activity [40,56–59]. This highlights a potential molecular mechanism for how the R1021C mutation can lead to increased lipid kinase activity.

**Figure 2.**
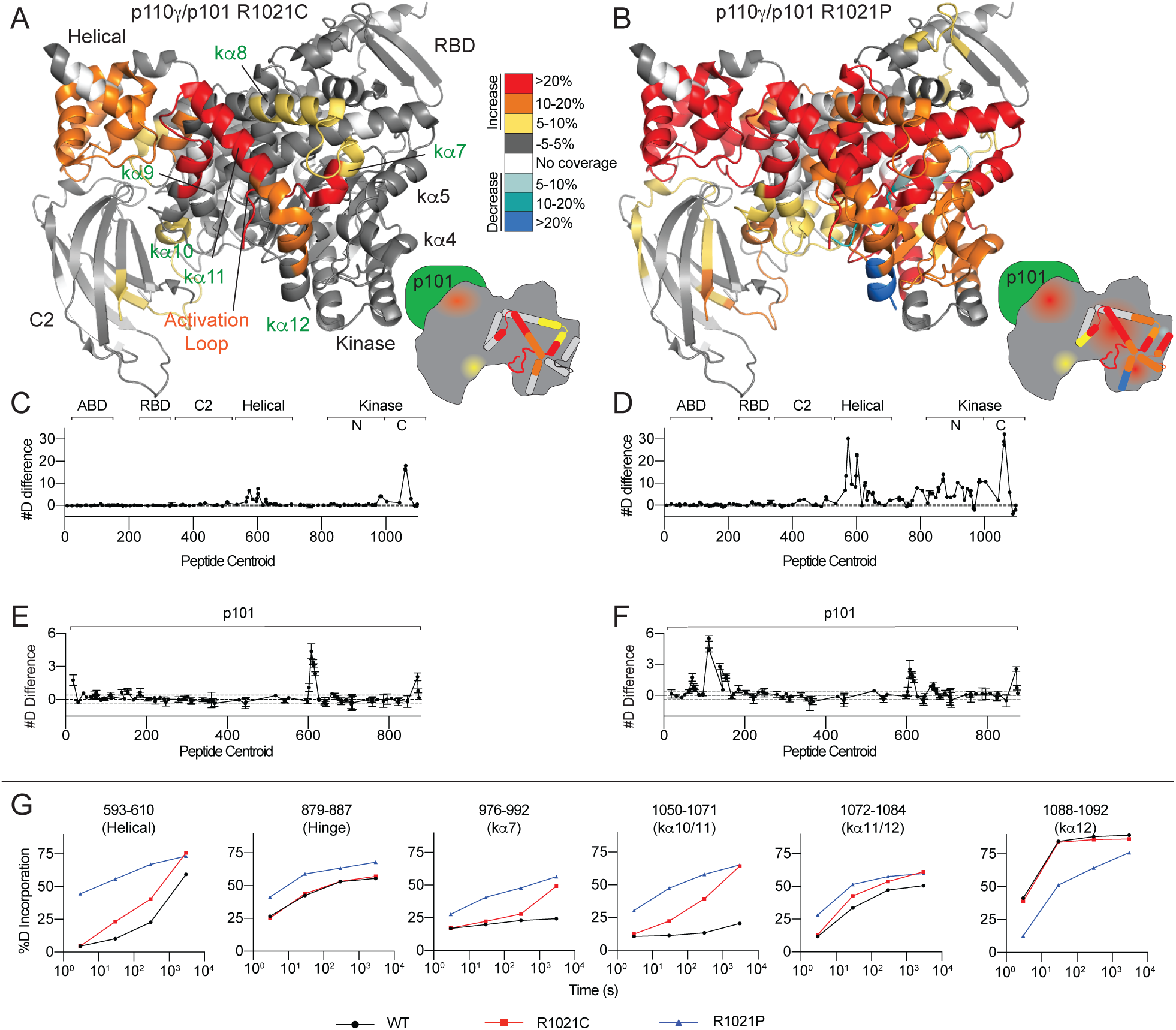
R1021C and R1021P mutations in p110γ are destabilising, with R1021P leading to global destabilization and R1021C leading to localised disruption of the C-terminal regulatory W1080 Tryptophan ‘lock’. **A+B.** Peptides showing significant deuterium exchange differences (>5 %, >0.4 kDa and p<0.01 in an unpaired two-tailed t-test) between wild-type and R1021C (A) and wild-type and R1021P (B) p110*γ*/p101 complexes are coloured on a model of p110*γ* (PDB: 6AUD)[54]. Differences in exchange are coloured according to the legend. **C+D.** The number of deuteron difference for the R1021C and R1021P mutants for all peptides analysed over the entire deuterium exchange time course for p110*γ*. Every point represents the central residue of an individual peptide. The domain location is noted above the primary sequence. A cartoon model of the p110*γ* structural model is shown according to the legend in panels A+B. Error is shown as standard deviation (n=3). **E+F.** The number of deuteron difference for the R1021C and R1021P mutants for all peptides analysed over the entire deuterium exchange time course for p101. Every point represents the central residue of an individual peptide. Error is shown as standard deviation (n=3). **G.** Selected p110*γ* peptides that showed decreases and increases in exchange are shown. The full list of all peptides and their deuterium incorporation is shown in supplementary data 1.

The R1021P mutation resulted in larger increases in exchange throughout almost the entire C2, helical, and kinase domains (Fig. 2D). Comparing the rates of hydrogen exchange between wild-type, R1021C, and R1021P showed many regions where R1021C and R1021P both caused increased exchange. However, for the majority of these regions the R1021P led to increased exchange at early (3 s) and late timepoints (3000 s) of exchange, indicative that this mutation was leading to significant disruption of protein secondary structure (Fig. 2G). This large-scale destabilization throughout the protein may explain the low yield and decreased kinase activity. The two mutations in R1021C and R1021P both caused increased exchange in the p101 subunit. Peptides spanning 602-623, and 865-877 of p101 showed similar increases in exchange for both R1021C and R1021P, with R1021P also leading to increased exchange in a peptide nearer the N-terminus (102-122) (Fig. 2E+F, S3). As there is no structural model for the p101 subunit, it is hard to unambiguously interpret this data, however, as these may represent increased exchange due to partial destabilization of the complex, our work provides initial insight into the p110*γ* contact site on p101.

### Molecular dynamics of p110γ R1021C and R1021P mutants

We carried out Gaussian-accelerated Molecular Dynamics (GaMD) simulations of wild-type p110*γ* and its R1021C and R1021P variants to provide additional insight into the underlying molecular mechanisms of how these mutations alter lipid kinase activity. Using the crystallographic structure of p110*γ* lacking the N-terminus [amino acids 144-1102, PDB: 6AUD [54]], we generated the activation loop and other neighboring loops as described in the methods, removed the co-crystallized ligand, and mutated R1021 to a cysteine and proline, resulting in three systems: WT, R1021C, and R1021P. Three replicas of fully solvated all-atom GaMD simulations were run for each model with AMBER18 achieving a cumulative extensive sampling of ∼3, ∼4.1, and ∼1.5 µs for WT, R1021C, and R1021P, respectively (Fig. 3A+B).

**Figure 3.**
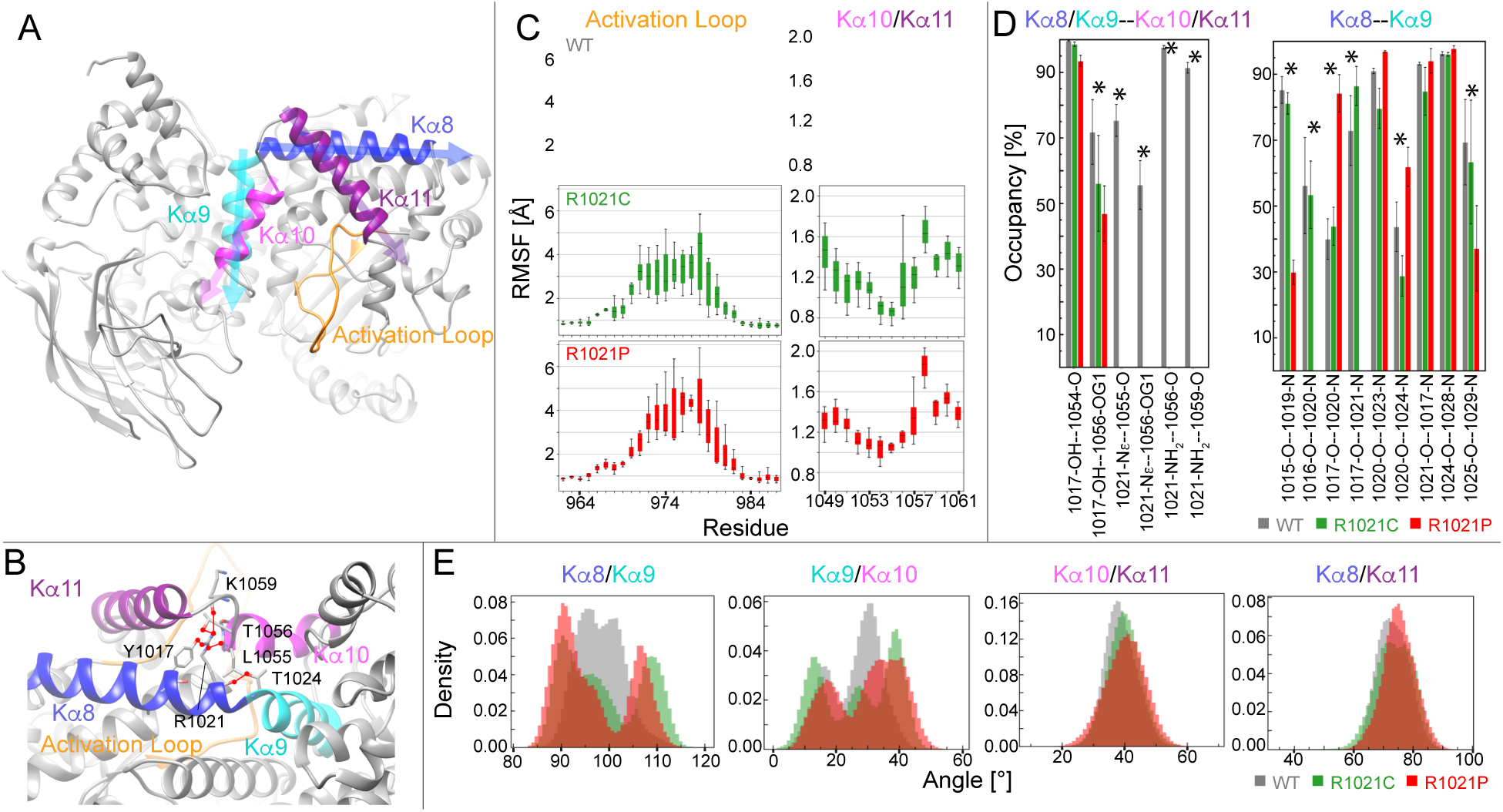
Molecular dynamics reveal that the R1021C and R1021P mutations show increased instability in p110γ. **A.** Model of p110*γ* showcasing the regulatory domain’s k*α*8 (995-1023), k*α*9 (1024-1037), k*α*10 (1045-1054), and k*α*11 (1057-1078) helices, and the activation loop (962-988). **B.** A zoomed-in snapshot of R1021 microenvironment showing residues in licorice. Hydrogen bonds with R1021 are drawn as red lines. **C.** RMSF [Å] of each residue’s C*α* and C*β* atoms in the activation loop and the k*α*10/k*α*11 helices, respectively. RMSF values for each atom across replicates are shown as a quantile plot, with error shown as standard deviation (n=3). **D.** The mean and standard deviation of hydrogen bond occupancies between the indicated helices/sets of helices across replicates (n=3). Asterisks indicate significant differences in occupancies. **E.** Inter-angle density distributions across all replicas between k*α*8, k*α*9, k*α*10, and k*α*11. In all panels, WT, R1021C, and R1021P are colored in grey, green, and red, respectively.

To quantify the effect of mutations on the structural dynamics of p110*γ*, we calculated the root-mean-square-fluctuation (RMSF) of residues neighboring the mutation site. RMSF was calculated to determine average flexibility of each residue’s C*α* and C*β* atoms around their mean position (Fig. 3C). This revealed increased fluctuations in the residues forming the loop between k*α*10 and k*α*11 in the mutated systems, specifically residues T1056, V1057, and G1058 at the C-terminus of k*α*10. Many of these residues participate in hydrogen bonds with R1021 in WT (Fig. 3B).

Analysis of the simulations revealed that mutation of R1021 results in disruption of the hydrogen bonding network between R1021 and L1055, T1056, and K1059 in the k*α*10-k*α*11 region. There were also alterations in the intra and inter helix hydrogen bonds in k*α*8, k*α*9, k*α*10, and k*α*11 (Fig. 3D, S4). Hydrogen bonding occupancies between Y1017 and T1056 decreased from 71% in WT to 56% and 45% in the R1021C and R1021P systems, respectively. Examining the k*α*8-k*α*9 backbone hydrogen bonding at the site of mutation, both mutations showed a disruption between C/P1021 and T1024. Additionally, the proline mutation showed complete disruption of backbone hydrogen bonds at A1016-L1020 and Y1017-P1021, decreased bonding occupancy at K1015-A1019 and N1025-I1029, and increased bonding occupancy of Y1017-L1020 and P1021-T1024. The notable increase in hydrogen bonding disruption in the R1021P compared to R1021C could be responsible for the increased destabilization observed by HDX-MS.

To obtain further insights into the dynamic behavior of the C-terminus of the kinase domain and how mutation of R1021 alters conformational dynamics, we monitored the fluctuations of four different angles formed between k*α*8, k*α*9, k*α*10, and k*α*11 (Fig. 3E). The simulations revealed increased angle fluctuations in the mutant simulations between k*α*8 and k*α*9, and k*α*9 and k*α*10, with a bimodal distribution in the k*α*8/k*α*9 angle compared to WT. There was also increased fluctuations in the activation loop in the mutants compared to WT (Figure 3C, Fig. S4).

### Multiple PI3Kγ inhibitors lead to allosteric conformational changes

Many of the differences in conformational dynamics observed by HDX-MS for the p110*γ* mutants were similar to previously observed allosteric changes caused by cyclopropyl ethyl containing isoindolinone compounds [34]. We performed HDX-MS experiments with seven potent PI3K inhibitors on free p110*γ* to define the role of allostery in PI3K*γ* inhibition. We analysed inhibitors that were selective for PI3K*γ* [AS-604850 [9], AZ2 [34], NVS-PI3-4 [15, 60], and IPI-549 [53]) as well as pan-PI3K inhibitors [PIK90 [61], Omipalisib [62], and Gedatolisib [63]]. Of these compounds only AS-604850, PIK90, and Omipalisib have been structurally characterized bound to p110*γ*. A table summarizing these compounds and their selectivity for different PI3K isoforms is shown in table S2. Deuterium exchange experiments were carried out with monomeric p110*γ* over 4 timepoints of deuterium exchange (3,30,300, and 3000 s). We obtained 180 peptides covering ∼89% percent of the exchangeable amides (Table S1). To verify that results on the free p110*γ* complex are relevant to the physiological p110*γ*/p101 complex, we also carried out experiments with the p110*γ*/p101 complex with Gedatolisib and IPI-549, with the free p110*γ* showing almost exactly the same differences as seen for the p110*γ*/p101 complex (Fig. S5).

Based on the H/D exchange differences observed with inhibitors present, we were able to classify the inhibitors into three broad groups. The first group contains the isoquinolinone compound IPI-549, the imidazo[1,2-*c*]quinazoline molecule PIK-90 and the thiazolidinedione compound AS-604850 (Fig. 4A+B). These compounds caused decreased exchange near the active site, with the primary region being protected being the hinge region between the N-and C-lobes of the kinase domain (Fig. 4B+E). No (IPI-549, AS-604850) or very small (PIK-90) increases in deuterium incorporation were observed (Fig 4A, S6), suggesting that there are limited large scale allosteric conformational changes for these compounds.

**Figure 4.**
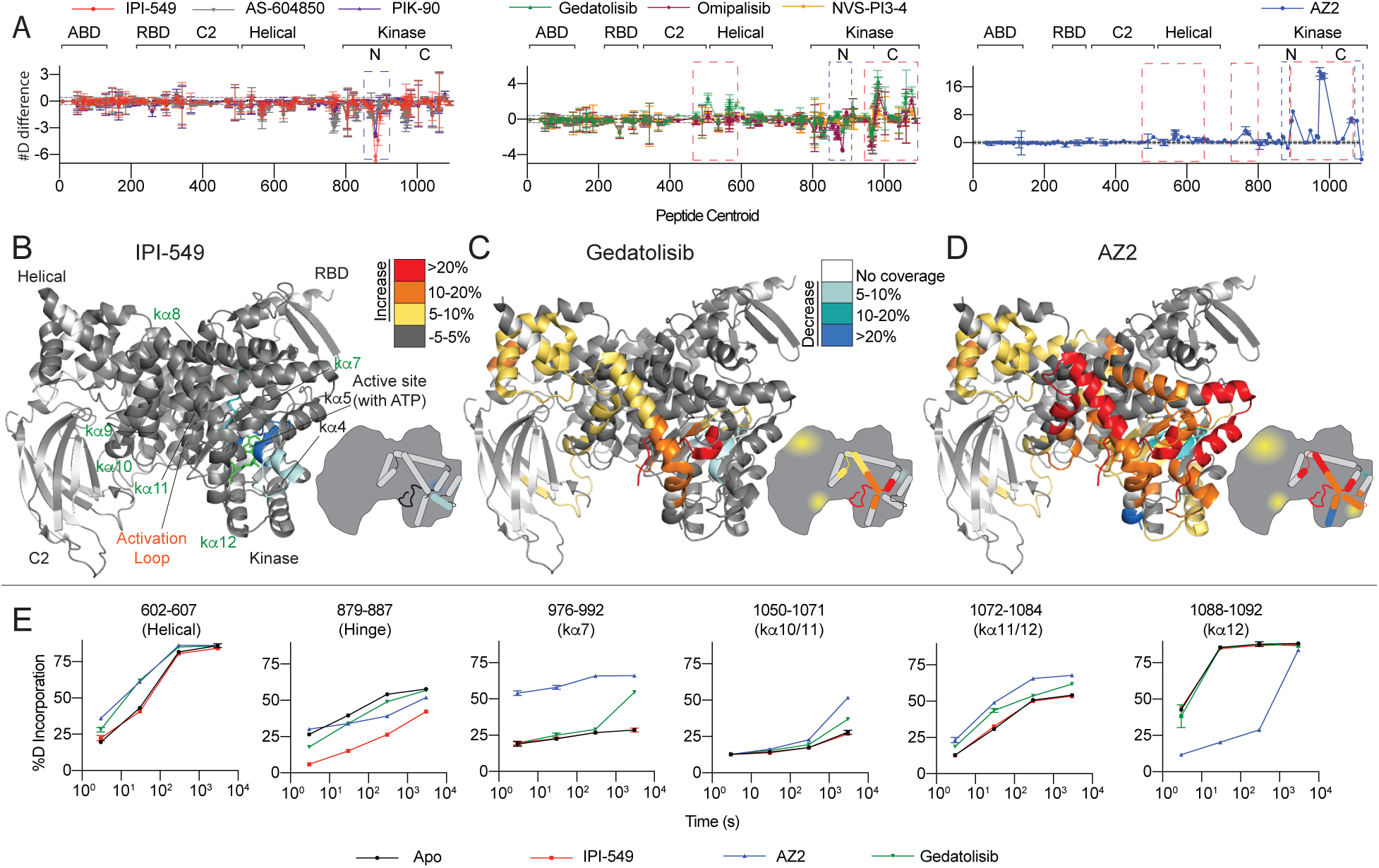
HDX-MS reveals that different classes of PI3K inhibitors lead to unique allosteric conformational changes. **A.** The number of deuteron difference for the 7 different inhibitors analysed over the entire deuterium exchange time course for p110*γ*. Every point represents the central residue of an individual peptide. The domain location is noted above the primary sequence. Error is shown as standard deviation (n=3). **B-D.** Peptides showing significant deuterium exchange differences (>5%, >0.4 kDa and p<0.01 in an unpaired two-tailed t-test) between wild-type and IPI-549 (**B**), Gedatolisib (**C**), and AZ2 (**D**) are coloured on a model of p110*γ* (PDB: 6AUD). Differences in exchange are mapped according to the legend. A cartoon model in the same format as Fig. 1 is shown as a reference. **E.** Selected p110*γ* peptides that showed decreases and increases in exchange are shown. The full list of all peptides and their deuterium incorporation is shown in supplementary data 1.

The H/D exchange experiments revealed a second class of inhibitors that showed decreased exchange at the active site, but also significant increases in exchange in the kinase and helical domains (Fig. 4A+C, S6). The second group includes the bis-morpholinotriazine molecule Gedatolisib, difluoro-benzene sulfonamide compound Omipalisib and the PI3K*γ*-specific thiazole derivative NVS-PI3-4. Binding of these inhibitors caused increased exchange in the helical domain, and multiple regions of the kinase regulatory motif, including k*α*7, k*α*10, k*α*11 and k*α*12. The peptide covering k*α*7 also spans the C-terminal end of the activation loop. Intriguingly, for the Gedatolisib molecule, the differences in H/D exchange matched very closely to those observed in the R1021C mutant. This suggests that the conformational changes induced by these compounds mimic the partially activated state that occurs in the R1021C mutant.

Finally, AZ2 caused large scale increased exposure throughout large regions of the helical and kinase domains (Fig. 4A+D), consistent with previous reports [34]. The same regulatory motif regions that showed increased exchange with Gedatolisib showed much larger changes with AZ2. Importantly, increased exchange was observed at earlier timepoints for AZ2 compared to Gedatolisib (example peptide 976-992 covering the activation loop and k*α*7), suggesting that AZ2 leads to a complete disruption of secondary structure, with Gedatolisib likely causing increased secondary structure dynamics (Fig. 4E).

This shows that multiple PI3K inhibitors can cause large scale allosteric conformational changes upon inhibitor binding, however, deciphering the molecular mechanism of these changes were hindered by lack of high-resolution structural information for many of these compounds.

### Structures of PI3Kγ bound to IPI-549, Gedatolisib, and NVS-PI3-4

To further define the molecular basis for how different inhibitors lead to allosteric conformational changes we solved the crystal structure of p110*γ* bound to IPI-549, Gedatolisib, and NVS-PI3-4 at resolutions of 2.55Å, 2.65Å, and 3.15Å, respectively (Fig. 5A-C, S6, S8). The inhibitor binding mode for all were unambiguous (Fig. S8).

**Fig 5:**
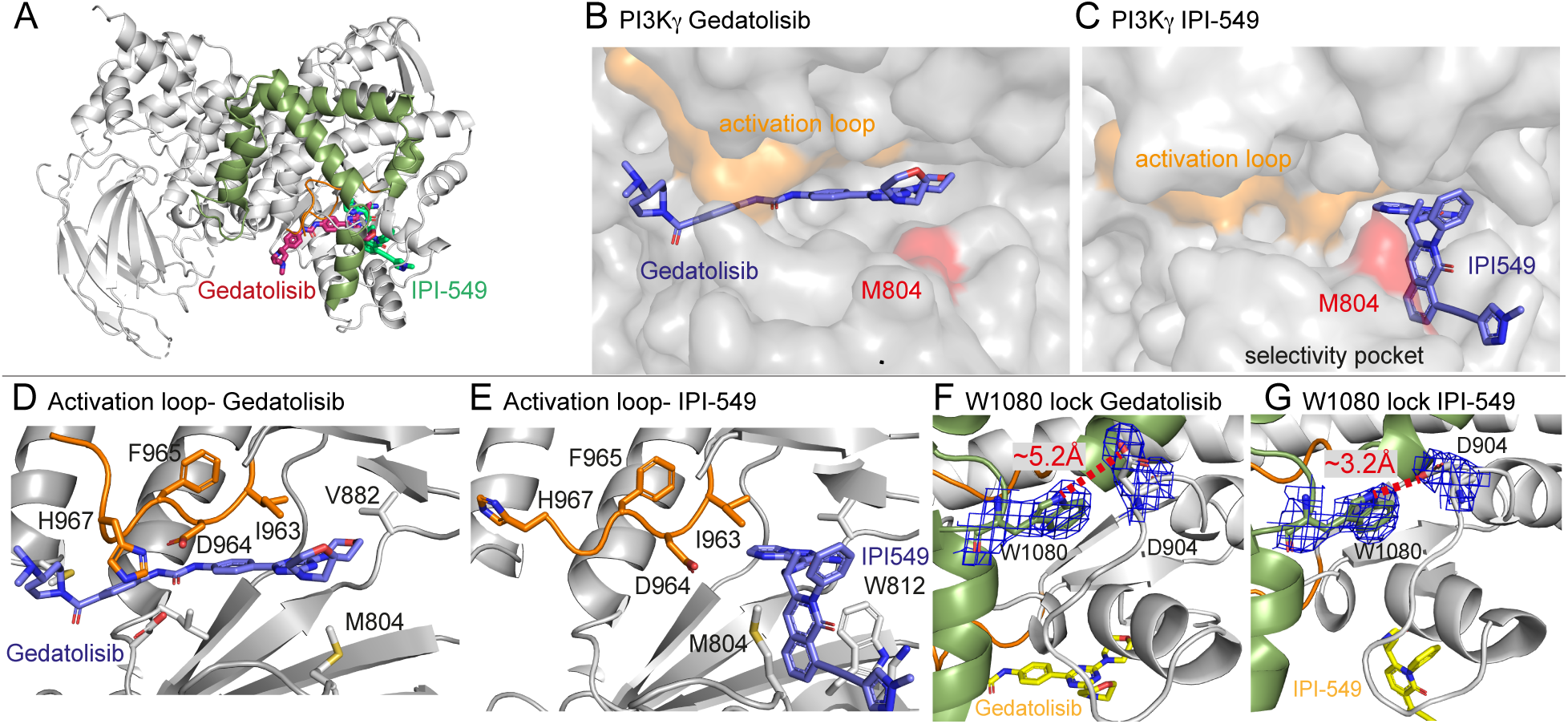
Structures of Gedatolisib and IPI-549 bound to p110γ. **A.** Overall structure of Gedatolisib (red) and IPI-549 (green) bound to p110*γ*. **B-C.** Comparison of Gedatolisib and IPI-549 bound to p110*γ* with the activation loop and selectivity pocket highlighted. M804 that changes conformation upon selectivity pocket opening is coloured red. **D-E.** Comparison of the conformation of the activation loop (orange) of p110*γ* when Gedatolisib or IPI-549 are bound, with residues in the activation loop labelled, specifically D964 and F965 of the DFG motif labelled. **F-G.** The Trp lock composed of W0180 is partially disrupted in the Gedatolisib structure compared to the IPI-549 structure. The interaction between W1080 and D904 is shown, with the distance between the two shown on each structure. The electron density from a feature enhanced map [65] around W1080 and D904 in each structure is contoured at 1.5 sigma.

These structures revealed insight into how IPI-549 and NVS-PI3-4 can achieve selectivity for PI3K*γ* (Fig. S7). All inhibitors formed the critical hydrogen bond with the amide hydrogen of V882 in the hinge, which is a conserved feature of ATP competitive PI3K kinase inhibitors. NVS-PI3-4 leads to opening of a p110*γ* unique pocket mediated by a conformational change in K883 (Fig. S7D-H). The opening of K883 is accommodated by it rotating into contact with D884 and T955. This opening would not be possible in p110*α* and p110*δ* as the corresponding K883 residue (L829 in p110*δ* and R852 in p110*α*) would clash with the corresponding T955 residue (R902 in p110*δ* and K924 in p110*α*) (Fig S7I-J). IPI-549 binds with a characteristic propeller shape, as seen for multiple p110*γ* and p110*δ* selective inhibitors [64]. IPI-549 leads to a conformational change in the orientation of M804, which opens the specificity pocket, primarily composed of W812 and M804 (Fig. 5C, S7). Comparison of IPI-549 bound to p110*γ* to the selective inhibitor Idelalisib bound to p110*δ* revealed a potential molecular mechanism for p110*γ* selectivity. Structure activity analysis of IPI-549 and its derivatives showed a critical role for substitutions at the alkyne position in achieving p110*γ* specificity[53]. The *N-*methylpyrazole group in IPI-549 projects out of the specificity pocket towards the k*α*1-k*α*2 loop. This loop is significantly shorter in p110*δ*, with a potential clash with bulkier alkyne derivatives (Fig. S7K-L). However, this cannot be the main driver of specificity, as a phenyl substituent of the alkyne had decreased selectivity of p110*γ* over p110*δ*, with hydrophilic heterocycles in this position being critical in p110*γ* selectivity[53]. A major difference in this pocket between p110*γ* and p110*δ* is K802 in p110*γ* (T750 in p110*δ*), with this residue making a pi-stacking interaction with W812. The *N-*methylpyrazole group packs against K802, with a bulkier group in this position likely to disrupt the pi stacking interaction, explaining the decreased potency for these compounds[53].

One of the most striking differences between the structure of Gedatolisib and IPI-549 bound to p110*γ* is the conformation of the N-terminus of the activation loop, including the residues that make up the DFG motif (Fig. 5B, D+E, S8). The majority of the activation loop is disordered in PI3K*γ* crystal structures, with the last residue being between 967 and 969. Gedatolisib makes extensive contacts with the activation loop, with H967 immediately following the DFG motif in a completely altered conformation. The interaction of the cyclopropyl motif in AZ2 with the activation loop has previously been proposed to be critical in mediating allosteric conformational changes. In addition to the change in the activation loop, there was a minor perturbation of the W1080 lock, with the Gedatolisib structure revealing a disruption of the hydrogen bond between W1080 and D904, with this bond maintained in the IPI-549 structure (Fig. 5F+G). The C-terminus of the activation loop and k*α*7 immediately following showed some of the largest changes upon inhibitor binding in HDX experiments. The k*α*7 helix is directly in contact with W1080, and we postulated that the conformational changes induced in the N-terminus of the activation loop may mediate these changes.

### Conformational selective inhibitors show altered specificity towards activating PI3Kγ mutant

We observed that HDX differences occurring in the R1021C mutant, were very similar to conformational changes observed for p110*γ* bound to Gedatolisib, particularly for the peptide spanning 976-992 in the activation loop. As this region is directly adjacent to the inhibitor binding site, we postulated that there may be altered inhibitor binding for the R1021C mutant. We carried out IC_50_ measurement for wild-type and R1021C p110*γ*/p101 with both IPI-549 and Gedatolisib (Fig. 6A). Gedatolisib was roughly three-fold more potent for the R1021C mutant over the wild-type, with no significant difference in IC_50_ values for IPI-549 compared to wild-type. This provides initial insight into how understanding the dynamics of activating mutations and inhibitors may be useful as a novel strategy towards designing mutant specific inhibitors.

**Fig. 6.**
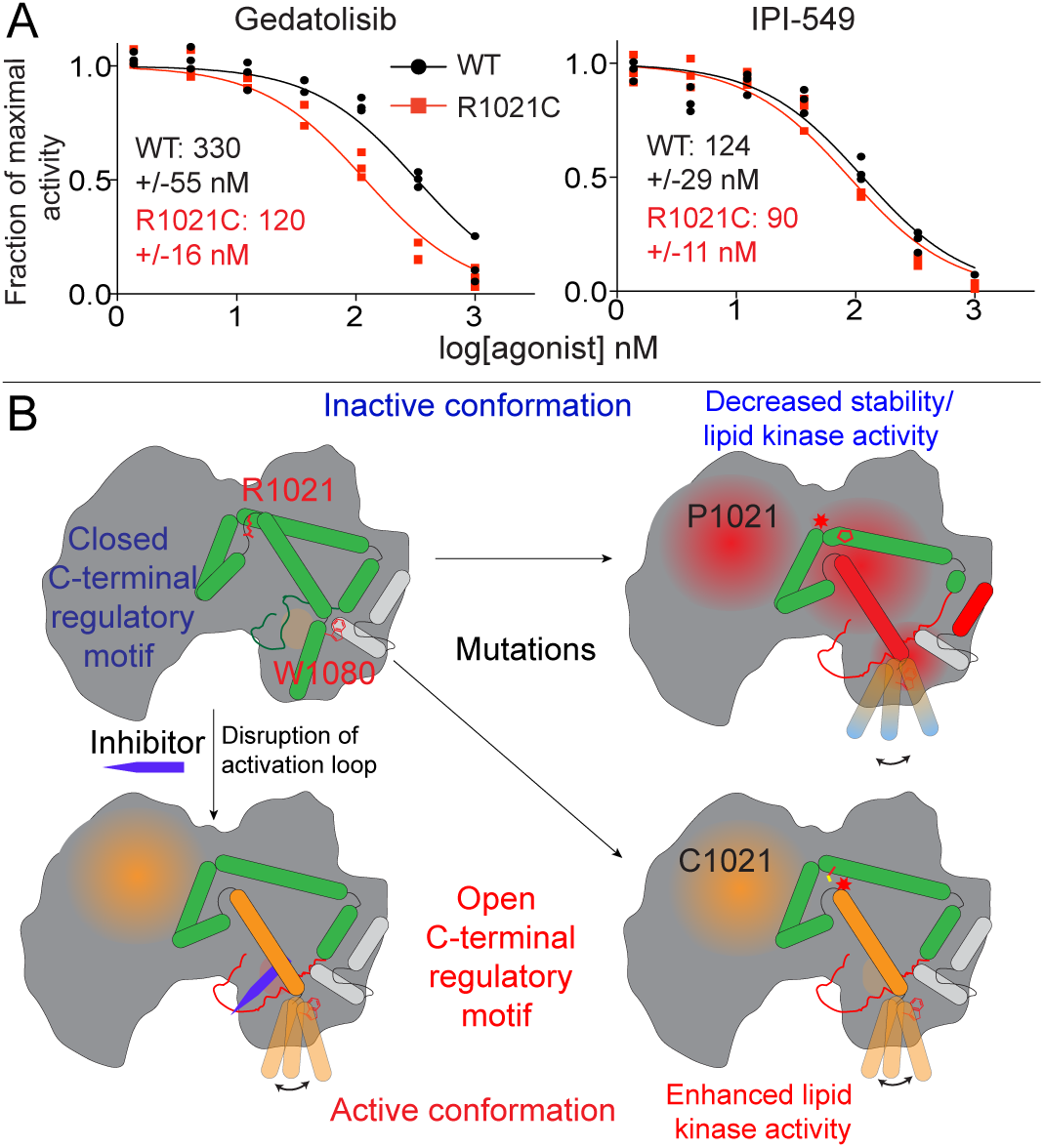
Activating mutations show slight differences in inhibition by allosteric inhibitors and model of PI3Kγ regulation. **A.** IC_50_ curves for wild-type and R1021C p110*γ*/p101 complexes. Assays were carried out with 5% C8 PIP_2_ / 95% PS vesicles at a final concentration of 1 mg/ml in the presence of 100 μM ATP and 1.5 μM lipidated G*βγ*. PI3K*γ* concentration was 4 nM for R1021C and 8 nM for WT. Error is shown as standard deviation (n=3) **B.** Model of conformational changes that occur upon mutation of the C-terminal motif and binding of activation loop interacting conformation selective inhibitors.

## Discussion

Understanding the molecular determinants of how mutations in PI3Ks lead to altered signalling in disease is vital in the design of targeted therapeutic strategies. The PI3K*γ* isoform is critical to maintain immune system function, and plays important roles in the regulation of the tumour microenvironment [7, 66]. Bi-allelic loss of function mutations in PI3K*γ* are a driver of human immunodeficiencies, and multiple inactivating mutations located in the C-terminal regulatory motif of the kinase domain have been described [46, 47]. Initial results linking deletion of PI3K*γ* to the development of colon cancer [67] have been disputed [68], and recent studies suggest that tumour growth and metastasis is attenuated in PI3K*γ* deficient mice [30, 69] and IPI-549 treated animals [29]. Inhibiting PI3K*γ* has shown promise as an immunomodulatory agent in generating an anti-tumour immune response [29, 30]. There have also been numerous reports of overexpression and single nucleotide variants in PIK3CG linked to cancer development in multiple tissues [69–76]. Oncogenic mutations in PIK3CG are widely distributed, which is distinct from the oncogenic hotspot mutations seen in the helical and kinase domain of PIK3CA. There has been limited analysis of the functional consequences of oncogenic PIK3CG mutants, with the R1021 residue in the regulatory motif of the kinase domain being unique, as mutations of this residue exist in both immunodeficiencies and tumours.

Here, we have described the biochemical and biophysical characterisation of both activating and inactivating disease linked R1021 mutations in the regulatory motif of the PI3K*γ* kinase domain. This has revealed that mutation of R1021 can lead to either kinase activation or inactivation. The R1021 in the k*α*8 helix is conserved across all class I PI3Ks, with it making a number of hydrogen bonds with residues in k*α*10 and k*α*11. Both R1021P and R1021C would lead to disruption of the hydrogen bonds with k*α*10 and k*α*11, however R1021P would also lead to disruption of the k*α*8 helix due to the altered dynamics introduced by the proline residue. HDX-MS results were consistent with this hypothesis, with R1021P leading to large scale conformational changes across the entire protein, with the main disruptions occurring in the helical and kinase domain. Molecular dynamics simulations revealed alterations in the fluctuation of the helices in the C-terminal regulatory motif for R1021P. The k*α*10 helix in the kinase domain extensively contacts the helical domain, with the altered orientation of this helix potentially revealing a mechanism of increased exchange in the helical domain. The R1021P mutation greatly destabilized the protein, with purification yields being >20-fold lower than wild-type, consistent with greatly decreased p110*γ* and p101 levels in patient T cells [46]. Consistent with previous reports we found greatly decreased lipid kinase activity for R1021P, although the enzyme maintained catalytic ability, as it showed greatly increased basal ATPase activity, which is similar to what occurs upon mutation of the W1080 lock or removal of the k*α*12 helix [34, 46]. This suggests a mechanism whereby R1021P mutation leads to large scale destabilization, and locks the enzyme into a lipid kinase inactive form.

The R1021C mutation in contrast, had enhanced lipid kinase activity, both basally, and upon G*βγ* activation. Increased conformational changes for this mutation were primarily localised to the C-terminal regulatory motif, with additional increased exchange occurring in the helical domain, although not to the same extent as seen in R1021P. Many of these changes in the C-terminal regulatory motif have been previously observed upon membrane binding [21], as well as upon binding to conformational selective inhibitors [34]. One of the largest changes in exchange occurred at the C-terminus of the activation loop and the beginning of k*α*7 which is in contact with the W1080 lock. We propose a model of how mutation of R1021 can lead to either activated or inactivated lipid kinase activity (Fig. 6B). The conformation of the C-terminal regulatory motif is critical in regulating lipid kinase activity, where minor perturbations (R1021C) can lead to disruption of multiple inhibitory contacts allowing for reorientation of the k*α*12 membrane binding helix and increased lipid kinase activity. For R1021P, this mutation leads to extensive conformational disruption throughout the protein, along with the C-terminal regulatory domain, which results in decreased protein stability and inactivation of kinase activity. Reinforcing this as a general mechanism important for class I PI3K regulation is that mutation of the equivalent R992 in PIK3CA to either Leu or Asn has been found in tumour samples [45].

This work corroborates the important role of the C-terminal regulatory motif in controlling PI3K lipid kinase activity. The orientation of this motif is critical in the regulation of all class I PI3Ks, although this is regulated by different molecular mechanisms in p110*α*, p110*β*, p110*δ*, and p110*γ*. The class IA PI3Ks require p85 regulatory subunits to stabilize the C-terminal regulatory motif, with the nSH2 of p85 interacting with and stabilising k*α*10 for all class IA PI3Ks [57, 77], and the cSH2 of p85 stabilising k*α*7, k*α*8, k*α*11 and k*α*12 for p110*β* and p110*δ* [59, 78]. The p110*γ* isoform is unique in that its C-terminal motif adopts an inhibited conformation in the absence of regulatory proteins. The C-terminal regulatory motif of p110*γ* can be post-translationally modified by phosphorylation of k*α*9 (T1024) by protein kinase A decreasing lipid kinase activity [79], while protein kinase C phosphorylates an adjacent area in the helical domain (S582) [80] increasing lipid kinase activity.

It has previously been noted that PI3K*γ* can be selectively targeted through a conformationally selective inhibitor, AZ2 [34]. This was mediated through a cyclopropyl moiety on AZ2, which putatively alters the orientation of the activation loop, leading to disruption of the inhibitory conformation of the C-terminal regulatory motif. Many of the changes observed for this inhibitor were similar to those seen in the R1021C and R1021P mutant. To interrogate if allosteric conformational changes were unique to cyclopropyl containing compounds, we screened a panel of pan-PI3K and PI3K*γ* selective inhibitors using HDX-MS. HDX-MS analysis of inhibitors bound to PI3K*γ* revealed distinct dynamics between compounds. The compounds PIK90, IPI549, and AS-604850 only caused decreased exchange at the active site. Comparison of the crystal structures of these compounds [9, 61] revealed similar conformation of the activation loop, with limited interaction between the inhibitors and the activation loop. AZ2, containing the cyclopropyl moiety led to large scale conformational changes consistent with previous results [34]. Intriguingly, the non-specific inhibitors Gedatolisib and Omipalisib caused increased exchange in many of the same regions that showed enhanced exchange with the R1021C mutant. Comparison of the crystal structures of these inhibitors [62] revealed more extensive interactions with the activation loop, and significant conformational rearrangement of the activation loop. Distinct from the AZ2 compound, neither Gedatolisib and Omipalisib show specificity for PI3K*γ* over class IA PI3Ks [62, 63]. Similar HDX-MS differences were observed for both the R1021C mutant and wild type bound to Gedatolisib. Gedatolisib showed increased potency versus R1021C over wild type PI3K*γ*, with a ∼3-fold decrease in IC50 values. Altogether, this suggests that R1021C induces a conformation similar to the wild type enzyme bound to Gedatolisib. This provides an intriguing approach for designing oncogenic PI3K specific inhibitors through further optimisation of the ATP competitive inhibitor moieties in the activation loop binding region.

Overall, this work provides novel insight into how the C-terminal regulatory motif of PI3K*γ* regulates lipid kinase activity, how oncogenic and immunodeficiency mutations can disrupt this regulation, and how we can exploit these conformational changes to develop isoform and mutant selective small molecule inhibitors. Further exploration of the dynamic regulation of the C-terminal regulatory motif of PI3Ks by mutations and inhibitors may reveal unique approaches to develop therapeutics for PI3K related human diseases.

## Methods

### Expression and Purification of PI3Kγ constructs

Full length monomeric p110*γ* (WT, R1021C) and p110*γ*/p101 complex (WT, R1021C, R1021P) were expressed in Sf9 insect cells using the baculovirus expression system. For the complex, the subunits were co-expressed from a MultiBac vector[81]. Following 55 hours of expression, cells were harvested by centrifuging at 1680 RCF (Eppendorf Centrifuge 5810 R) and the pellets were snap-frozen in liquid nitrogen. Both the monomer and the complex were purified identically through a combination of nickel affinity, streptavidin affinity and size exclusion chromatographic techniques.

Frozen insect cell pellets were resuspended in lysis buffer (20 mM Tris pH 8.0, 100 mM NaCl, 10 mM imidazole pH 8.0, 5% glycerol (v/v), 2 mM beta-mercaptoethanol (*β*ME), protease inhibitor (Protease Inhibitor Cocktail Set III, Sigma)) and sonicated for 2 minutes (15s on, 15s off, level 4.0, Misonix sonicator 3000). Triton-X was added to the lysate to a final concentration of 0.1% and clarified by spinning at 15,000 g for 45 minutes (Beckman Coulter JA-20 rotor). The supernatant was loaded onto a 5 mL HisTrap™ FF crude column (GE Healthcare) equilibrated in NiNTA A buffer (20 mM Tris pH 8.0, 100 mM NaCl, 20 mM imidazole pH 8.0, 5% (v/v) glycerol, 2 mM *β*ME). The column was washed with high salt NiNTA A buffer (20 mM Tris pH 8.0, 1 M NaCl, 20 mM imidazole pH 8.0, 5% (v/v) glycerol, 2 mM *β*ME), NiNTA A buffer, 6% NiNTA B buffer (20 mM Tris pH 8.0, 100 mM NaCl, 250 mM imidazole pH 8.0, 5% (v/v) glycerol, 2 mM *β*ME) and the protein was eluted with 100% NiNTA B. The eluent was loaded onto a 5 mL StrepTrap™ HP column (GE Healthcare) equilibrated in gel filtration buffer (20mM Tris pH 8.5, 100 mM NaCl, 50 mM Ammonium Sulfate and 0.5 mM tris(2-carboxyethyl) phosphine (TCEP)). The column was washed with the same buffer and loaded with tobacco etch virus protease. After cleavage on the column overnight, the protein was eluted in gel filtration buffer. The eluent was concentrated in a 50,000 MWCO Amicon Concentrator (Millipore) to <1 mL and injected onto a Superdex™ 200 10/300 GL Increase size-exclusion column (GE Healthcare) equilibrated in gel filtration buffer. After size exclusion, the protein was concentrated, aliquoted, frozen and stored at −80°C.

For crystallography, p110*γ* (144-1102) was expressed in Sf9 insect cells for 72 hours. The cell pellet was lysed and the lysate was subjected to nickel affinity purification as described above. The eluent was loaded onto HiTrap^TM^ Heparin HP cation exchange column equilibrated in Hep A buffer (20 mM Tris pH 8.0, 100 mM NaCl, 5% glycerol and 2 mM *β*ME). A gradient was started with Hep B buffer (20 mM Tris pH 8.0, 1 M NaCl, 5% glycerol and 2 mM *β*ME) and the fractions containing the peak were pooled. This was then loaded onto HiTrap^TM^ Q HP anion exchange column equilibrated with Hep A and again subjected to a gradient with Hep B. The peak fractions were pooled, concentrated on a 50,000 MWCO Amicon Concentrator (Millipore) to <1 mL and injected onto a Superdex™ 200 10/300 GL Increase size-exclusion column (GE Healthcare) equilibrated in gel filtration buffer (20 mM Tris pH 7.2, 0.5 mM (NH_4_)_2_SO_4_, 1% ethylene glycol, 0.02% CHAPS and 5 mM DTT). Protein from size exclusion was concentrated to >5 mg/mL, aliquoted, frozen and stored at −80°C.

### Expression and Purification of lipidated Gβγ

Full length, lipidated Gβ*γ* was expressed in Sf9 insect cells and purified as described previously[82]. After 65 hours of expression, cells were harvested and the pellets were frozen as described above. Pellets were resuspended in lysis buffer (20 mM HEPES pH 7.7, 100 mM NaCl, 10 mM *β*ME, protease inhibitor (Protease Inhibitor Cocktail Set III, Sigma)) and sonicated for 2 minutes (15s on, 15s off, level 4.0, Misonix sonicator 3000). The lysate was spun at 500 RCF (Eppendorf Centrifuge 5810 R) to remove intact cells and the supernatant was centrifuged again at 25,000 g for 1 hour (Beckman Coulter JA- 20 rotor). The pellet was resuspended in lysis buffer and sodium cholate was added to a final concentration of 1% and stirred at 4°C for 1 hour. The membrane extract was clarified by spinning at 10,000 g for 30 minutes (Beckman Coulter JA-20 rotor). The supernatant was diluted 3 times with NiNTA A buffer (20 mM HEPES pH 7.7, 100 mM NaCl, 10 mM Imidazole, 0.1% C12E10, 10mM *β*ME) and loaded onto a 5 mL HisTrap™ FF crude column (GE Healthcare) equilibrated in the same buffer. The column was washed with NiNTA A, 6% NiNTA B buffer (20 mM HEPES pH 7.7, 25 mM NaCl, 250 mM imidazole pH 8.0, 0.1% C12E10, 10 mM *β*ME) and the protein was eluted with 100% NiNTA B. The eluent was loaded onto HiTrap^TM^ Q HP anion exchange column equilibrated in Hep A buffer (20 mM Tris pH 8.0, 8 mM CHAPS, 2 mM Dithiothreitol (DTT)). A gradient was started with Hep B buffer (20 mM Tris pH 8.0, 500 mM NaCl, 8 mM CHAPS, 2 mM DTT) and the protein was eluted in ∼50% Hep B buffer. The eluent was concentrated in a 30,000 MWCO Amicon Concentrator (Millipore) to < 1 mL and injected onto a Superdex^TM^ 75 10/300 GL size exclusion column (GE Healthcare) equilibrated in Gel Filtration buffer (20 mM HEPES pH 7.7, 100 mM NaCl, 10 mM CHAPS, 2 mM TCEP). Fractions containing protein were pooled, concentrated, aliquoted, frozen and stored at −80°C.

### Expression and Purification of Lipidated HRas G12V

Full-length HRas G12V was expressed by infecting 500 mL of Sf9 cells with 5 mL of baculovirus. Cells were harvested after 55 hours of infection and frozen as described above. The frozen cell pellet was resuspended in lysis buffer (50 mM HEPES pH 7.5, 100 mM NaCl, 10 mM *β*ME and protease inhibitor (Protease Inhibitor Cocktail Set III, Sigma)) and sonicated on ice for 1 minute 30 seconds (15s ON, 15s OFF, power level 4.0) on the Misonix sonicator 3000. Triton-X 114 was added to the lysate to a final concentration of 1%, mixed for 10 minutes at 4°C and centrifuged at 25,000 rpm for 45 minutes (Beckman Ti-45 rotor). The supernatant was warmed to 37°C for few minutes until it turned cloudy following which it was centrifuged at 11,000 rpm at room temperature for 10 minutes (Beckman JA-20 rotor) to separate the soluble and detergent-enriched phases. The soluble phase was removed, and Triton-X 114 was added to the detergent-enriched phase to a final concentration of 1%. Phase separation was performed 3 times. Imidazole pH 8.0 was added to the detergent phase to a final concentration of 15 mM and the mixture was incubated with Ni-NTA agarose beads (Qiagen) for 1 hour at 4°C. The beads were washed with 5 column volumes of Ras-NiNTA buffer A (20mM Tris pH 8.0, 100mM NaCl, 15mM imidazole pH 8.0, 10mM *β*ME and 0.5% Sodium Cholate) and the protein was eluted with 2 column volumes of Ras-NiNTA buffer B (20mM Tris pH 8.0, 100mM NaCl, 250mM imidazole pH 8.0, 10mM *β*ME and 0.5% Sodium Cholate). The protein was buffer exchanged to Ras-NiNTA buffer A using a 10,000 kDa MWCO Amicon concentrator, where protein was concentrated to ∼1mL and topped up to 15 mL with Ras- NiNTA buffer A and this was repeated a total of 3 times. GTP*γ*S was added in 2-fold molar excess relative to HRas along with 25 mM EDTA. After incubating for an hour at room temperature, the protein was buffer exchanged with phosphatase buffer (32 mM Tris pH 8.0, 200 mM Ammonium Sulphate, 0.1 mM ZnCl2, 10 mM *β*ME and 0.5% Sodium Cholate). 1 unit of immobilized calf alkaline phosphatase (Sigma) was added per milligram of HRas along with 2-fold excess nucleotide and the mixture was incubated for 1 hour on ice. MgCl_2_ was added to a final concentration of 30 mM to lock the bound nucleotide. The immobilized phosphatase was removed using a 0.22-micron spin filter (EMD Millipore). The protein was concentrated to less than 1 mL and was injected onto a Superdex^TM^ 75 10/300 GL size exclusion column (GE Healthcare) equilibrated in gel filtration buffer (20 mM HEPES pH 7.7, 100 mM NaCl, 10 mM CHAPS, 1 mM MgCl2 and 2 mM TCEP). The protein was concentrated to 1 mg/mL using a 10,000 kDa MWCO Amicon concentrator, aliquoted, snap-frozen in liquid nitrogen and stored at −80°C.

### Lipid Vesicle Preparation

For kinase assays comparing WT and mutant activities, lipid vesicles containing 5% brain phosphatidylinositol 4,5- bisphosphate (PIP2), 20% brain phosphatidylserine (PS), 50% egg-yolk phosphatidylethanolamine (PE), 10% egg-yolk phosphatidylcholine (PC), 10% cholesterol and 5% egg-yolk sphingomyelin (SM) were prepared by mixing the lipids dissolved in organic solvent. The solvent was evaporated in a stream of argon following which the lipid film was desiccated in a vacuum for 45 minutes. The lipids were resuspended in lipid buffer (20 mM HEPES pH 7.0, 100 mM NaCl and 10 % glycerol) and the solution was sonicated for 15 minutes. The vesicles were subjected to five freeze thaw cycles and extruded 11 times through a 100-nm filter (T&T Scientific: TT-002-0010). The extruded vesicles were sonicated again for 5 minutes, aliquoted and stored at −80°C. For inhibitor response assays, lipid vesicles containing 95% PS and 5% C8-PIP2 were used. PS was dried and desiccated as described above. The lipid film was mixed and resuspended with C8-PIP2 solution (2.5 mg/mL in lipid buffer). Following this, vesicles were essentially prepared the same way as described above. All vesicles were stored at 5 mg/mL.

### Lipid Kinase assays

All lipid kinase activity assays employed the Transcreener ADP2 Fluorescence Intensity (FI) Assay (Bellbrook labs) which measures ADP production. For assays comparing the activities of mutants, final concentrations of PM-mimic vesicles were 1 mg/mL, ATP was 100 μM ATP and lipidated Gβ*γ*/HRas were at 1.5 μM. 2 μL of a PI3K solution at 2X final concentration was mixed with 2 μL substrate solution containing ATP, vesicles and Gβ*γ*/HRas or Gβ*γ*/HRas gel filtration buffer and the reaction was allowed to proceed for 60 minutes at 20°C. The reaction was stopped with 4 μL of 2X stop and detect solution containing Stop and Detect buffer, 8 nM ADP Alexa Fluor 594 Tracer and 93.7 μg/mL ADP2 Antibody IRDye QC-1 and incubated for 50 minutes. The fluorescence intensity was measured using a SpectraMax M5 plate reader at excitation 590 nm and emission 620 nm. This data was normalized against a 0-100% ADP window made using conditions containing either 100 μM ATP/ADP with vesicles and kinase buffer. % ATP turnover was interpolated from an ATP standard curve obtained from performing the assay on 100 μM (total) ATP/ADP mixtures with increasing concentrations of ADP using Prism 7. All specific activities of lipid kinase activity were corrected for the basal ATPase activity by subtracting the specific activity of the WT/mutant protein in the absence of vesicles/activators.

For assays measuring inhibitor response, substrate solutions containing vesicles, ATP and Gβ*γ* at 4X final concentration (as described above) were mixed with 4X solutions of inhibitor dissolved in lipid buffer (<1% DMSO) in serial to obtain 2X substrate solutions with inhibitors at the various 2X concentrations. 2 μL of this solution was mixed with 2 μL of 2X protein solution to start the reaction and allowed to proceed for 60 minutes at 37 °C. Following this, the reaction was stopped and the intensity was measured. The raw data was normalized against a 0-100% ADP window as described above. The % inhibition was calculated by comparison to the activity with no inhibitor to obtain fraction activity remaining.

### Hydrogen Deuterium Exchange Mass Spectrometry (HDX-MS)

HDX experiments were performed similarly as described before [40]. For HDX with mutants, 3 μL containing 13 picomoles of protein was incubated with 8.25 μL of D_2_O buffer (20mM HEPES pH 7.5, 100 mM NaCl, 98% (v/v) D_2_O) for four different time periods (3, 30, 300, 3000 s at 20 °C). After the appropriate time, the reaction was stopped with 57.5 μL of ice-cold quench buffer (2M guanidine, 3% formic acid), immediately snap frozen in liquid nitrogen and stored at −80 °C. For HDX with inhibitors, 5 μL of p110*γ* or p110*γ*/p101 at 2 μM was mixed with 5 μL of inhibitor at 4 μM in 10% DMSO or 5 μL of blank solution containing 10% DMSO and incubated for 20 minutes on ice. 40 μL of D2O buffer was added to this solution to start the exchange reaction which was allowed to proceed for four different time periods (3, 30, 300, 3000 s at 20 °C). After the appropriate time, the reaction was terminated with 20 μL of ice-cold quench buffer and the samples were frozen.

Protein samples were rapidly thawed and injected onto an ultra-high pressure liquid chromatography (UPLC) system at 2 °C. Protein was run over two immobilized pepsin columns (Trajan, ProDx protease column, PDX.PP01-F32 and Applied Biosystems, Porosyme, 2-3131-00) at 10 °C and 2 °C at 200 μl/min for 3 min, and peptides were collected onto a VanGuard precolumn trap (Waters). The trap was subsequently eluted in line with an Acquity 1.7-μm particle, 100 × 1 mm2 C18 UPLC column (Waters), using a gradient of 5–36% B (buffer A, 0.1% formic acid; buffer B, 100% acetonitrile) over 16 min. Mass spectrometry experiments were performed on an Impact II TOF (Bruker) acquiring over a mass range from 150 to 2200 m/z using an electrospray ionization source operated at a temperature of 200 °C and a spray voltage of 4.5 kV. Peptides were identified using data-dependent acquisition methods following tandem MS/MS experiments (0.5-s precursor scan from 150–2000 m/z; 12 0.25-s fragment scans from 150–2000 m/z). MS/MS datasets were analysed using PEAKS7 (PEAKS), and a false discovery rate was set at 1% using a database of purified proteins and known contaminants.

HD-Examiner software (Sierra Analytics) was used to automatically calculate the level of deuterium incorporation into each peptide. All peptides were manually inspected for correct charge state and presence of overlapping peptides. Deuteration levels were calculated using the centroid of the experimental isotope clusters. The results for these proteins are presented as relative levels of deuterium incorporation, and the only control for back exchange was the level of deuterium present in the buffer (62% for experiments with mutants and 75.5% for experiments with inhibitors). Changes in any peptide at any time point greater than both 5% and 0.4 Da between conditions with a paired t test value of p < 0.01 were considered significant. The raw HDX data are shown in two different formats. The raw peptide deuterium incorporation graphs for a selection of peptides with significant differences are shown, with the raw data for all analyzed peptides in the source data. To allow for visualization of differences across all peptides, we utilized number of deuteron difference (#D) plots. These plots show the total difference in deuterium incorporation over the entire H/D exchange time course, with each point indicating a single peptide. The mass spectrometry proteomics data have been deposited to the ProteomeXchange Consortium via the PRIDE partner repository[83] with the dataset identifier PXD021132.

### X-ray crystallography

p110*γ* (144-1102) was crystallized from a grid of 2μl sitting drops at 1:1, 2:1 and 3:1 protein to reservoir ratios at 18°C. Protein at 4 mg/mL (in 20 mM Tris pH 7.2, 0.5 mM (NH_4_)_2_SO_4_, 1% ethylene glycol, 0.02% CHAPS and 5 mM DTT) was mixed with reservoir solution containing 100 mM Tris pH 7.5, 250 mM (NH_4_)_2_SO_4_ and 20-22% PEG 4000. Large multinucleate crystals were generated in these drops. Inhibitor stocks were prepared at concentrations of 0.01 mM, 0.1 mM and 1 mM in cryo-protectant solution containing 100 mM Tris pH 7.5, 250 mM (NH_4_)_2_SO_4_, 23% PEG 4000 and 14% glycerol. Inhibitors at increasing concentrations were added to the drops stepwise every 1 hour. After overnight incubation with the inhibitor, single crystals were manually broken from the multi-nucleates and soaked in a fresh drop containing 1 mM inhibitor in cryo-protectant before being immediately frozen in liquid nitrogen.

Diffraction data for PI3K*γ* crystals were collected on beamline 08ID-1 of the Canadian Light Source. Data was collected at 0.97949. Data were processed using XDS [84]. Phases were initially obtained by molecular replacement using Phaser [85] using PDB: 2CHW for the IPI-549 complex [61], and 5JHA for Gedatolisib and NVS-PI3-4 [86]. Iterative model building and refinement were performed in COOT [87] and phenix.refine [88]. Refinement was carried out with rigid body refinement, followed by translation/libration/screw B-factor and xyz refinement. The final model was verified in Molprobity [89] to examine all Ramachandran and Rotamer outliers. Data collection and refinement statistics are shown in Table S3. The crystallography data has been deposited in the protein data bank with accession numbers (PDB: 7JWE, 7JX0, 7JWZ).

### Molecular Dynamics: Missing loops modelling

The employed crystallographic structures of the p110*γ* protein reveal several missing gaps corresponding to flexible loops within range of the ligand-binding site: the activation loop (residues 968-981), and loops connecting the C2 and helical domains (residues 435-460 and 489-497). These missing gaps were modelled as disordered loops using Modeller9.19 [90]. Keeping the crystallographic coordinates fixed, 50 models were independently generated for each system. The wild type (WT), R1021C, and R1021P systems used PDB ID 6AUD [54] with their corresponding mutations in the mutant systems. The alignment used by Modeller between the crystallographic structure sequences and the FASTA sequence of p110*γ* (Uniprot ID P48736) were generated using Clustal Omega [91]. The top models were visually inspected to discard those in which loops were entangled in a knot or clashed with the rest of the structure. Lastly, from the remaining models, three were selected for each system to initiate simulations in triplicates.

### Molecular Dynamics: System preparation

The generated models were prepared using tleap program of the AMBER package [92]. The systems were parametrized using the general AMBER force field (GAFF) using ff14sb for the protein [93]. The systems were fully solvated with explicit water molecules described using the TIP3P model [94], adding K+ and Cl- counterions to neutralize the total charge. The total number of atoms is 97,861 for WT (size: 116 Å × 95 Å × 94 Å), 100,079 for R1021C (size: 116 Å × 95 Å × 94 Å), 97,861 for R1021P (size: 116 Å × 95 Å × 94 Å).

### Gaussian accelerated Molecular Dynamics (GaMD)

All-atom MD simulations were conducted using the GPU version of AMBER18 [92]. The systems were initially relaxed through a series of minimization, heating, and equilibration cycles. During the first cycle, the protein was restrained using a harmonic potential with a force constant of 10 kcal/mol-Å^2^, while the solvent, and ions were subjected to an initial minimization of 2000 steps using the steepest descent approach for 1000 steps and conjugate gradient approach for another 1000 steps. The full system (protein + solvent) was then similarly minimized for 1000 and 4000 steps using the steepest descent and conjugate gradient approaches, respectively. Subsequently, the temperature was incrementally changed from 100 to 300 K for 10 ps at 2 fs/step (NVT ensemble). Next, the systems were equilibrated for 200 ps at 1 atom and 300K (NPT ensemble), and for 200ps at 300K (NVT ensemble). Lastly, more equilibration simulations were run in the NVT ensemble in two steps; all systems were simulated using conventional MD for 50 ns and GaMD for 50ns. Temperature control (300 K) and pressure control (1 atm) were performed via Langevin dynamics and Berendsen barostat, respectively. Production GaMD were simulated for ∼3 µs for WT, ∼4.1 µs R1021C, ∼1.5 µs for R1021P. GaMD is an unconstrained enhanced sampling approach that works by adding a harmonic boost potential to smooth biomolecular potential energy surface and reduce the system energy barriers [95]. Details of the GaMD method have been extensively described in previous studies [95, 96].

### GaMD analysis: Principal component analysis (PCA)

PCA was performed using the sklearn.decomposition.PCA function in the *Scikit-learn* library using python3.6.9. First, all simulations were aligned with *mdtraj* [97] onto the same initial coordinates using C*α* atoms of the kinase domain (residues 726–1088). Next, simulation coordinates of each domain of interest (for example k*α*9-k*α*10) from all systems (WT, R1021C, and R1021P) and replicas were concatenated and used to fit the transformation function. Subsequently, the fitted transformation function was applied to reduce the dimensionality of each domain’s simulation C*α* coordinates. It is important to note that all systems are transformed into the same PC space to evaluate the simulation variance across systems.

### GaMD analysis: Angles calculation

Inter-helical angles were calculated using in-house python scripts along with *mdtraj* [97] as the angle between two vectors representing the principal axis along each helix. Each principal axis connects two points corresponding to the center of mass (COM) of the first and last turn from each helix. For k*α*8, points 1 and 2 are represented by the COM of residues 1020-1023 and 1004-1007 C*α* coordinates, respectively. For k*α*9, points 1 and 2 are represented by the COM of residues 1024-1027 and 1034-1037 C*α* coordinates, respectively. For k*α*10, points 1 and 2 are represented by the COM of residues 1053- 1056 and 1046-1049 C*α* coordinates, respectively. For k*α*11, points 1 and 2 are represented by the COM of residues 1062-1065 and 1074-1077 C*α* coordinates, respectively. Angles were computed at each frame along the trajectories after structural alignment onto the initial coordinates using the C*α* atoms of the kinase domain (residues 726–1088) as a reference.

### GaMD analysis: Hydrogen bonds calculation

Hydrogen bonds were calculated using the *baker hubbard* command implemented with *mdtraj*[97] Occupancy (%) was determined by counting the number of frames in which a specific hydrogen bond was formed with respect to the total number of frames and then averaged across replicas.

### GaMD analysis: Root-mean-square-fluctuations (RMSF)

RMSF was calculated using in-house python scripts along with *mdtraj*[97] RMSF was computed for each residue atom and represented as box plot to show the range of RMSF values across replicas. The trajectories were aligned onto the initial coordinates using the Cα atoms of the kinase domain (residues 726–1088) as a reference.

### PI3K Inhibitors

Compounds were purchased from companies indicated below in ≥ 95% purity (typical 98% pure). IPI-549[53] was from ChemieTex (Indianapolis, USA, #CT-IPI549); PIK-90 [61] from Axon Medchem (Groningen, The Netherlands, #Axon1362); AS-604850 (PI 3-Kγ Inhibitor II, Calbiochem) [9] from Sigma Aldrich (#528108); Gedatolisib (PF-05212384, PKI587) [63] from Bionet (Camelford, UK, #FE-0013); Omipalisib (GSK2126458, GSK458) [62] from LuBioScience GmbH (Zurich, Switzerland, #S2658); NVS-PI3-4 [15, 60] and AZg1 (AZ2) [34] from Haoyuan Chemexpress Co., Ltd. (Shanghai, China, #HY-133907 and #HY-111570, respectively).

## Acknowledgements

J.E.B. is supported by a new investigator grant from the Canadian Institute of Health Research (CIHR), a Michael Smith Foundation for Health Research (MSFHR) Scholar award (17686), and an operating grant from the Cancer Research Society (CRS-24368). R.E.A and Z.G. are supported in part by NIH GM132826. M.P.W. is funded by the Stiftung für Krebsbekämpfung grant 341, the Swiss National Science Foundation grant 310030_189065, the Novartis Foundation for medical-biological Research grant 14B095; and the Innosuisse grant 37213.1 IP-LS. We appreciate feedback from members of the Burke lab during preparation.

## Data availability statement

The crystallography data has been deposited in the protein data bank with accession numbers (PDB: 7JWE, 7JX0, 7JWZ). The mass spectrometry proteomics data have been deposited to the ProteomeXchange Consortium via the PRIDE partner repository[83] with the dataset identifier PXD021132. All data generated or analyzed during this study are included in the manuscript and supporting files. Specifically biochemical kinase assay data are included in the source data files.

## Supplemental information: Supplemental Figures and Tables

**Figure S1 (relates to Fig 1).**
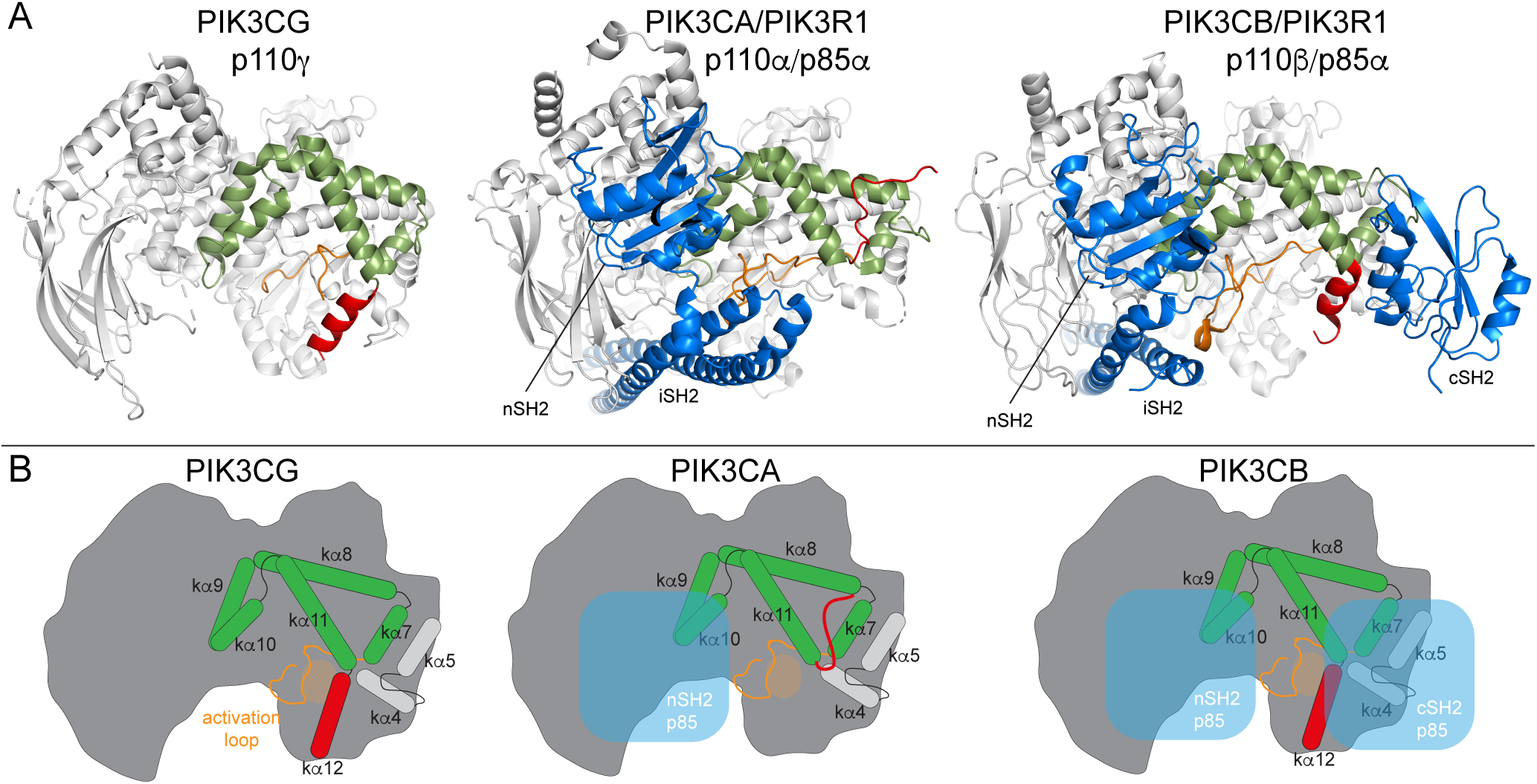
Comparing the different regulatory mechanisms that maintain the c-terminal regulatory motif in a inhibited state in the class I PI3Ks. **(A)** A structural model comparing the architecture of the C-terminal regulatory motif in PIK3CG (PI3K*γ*, PDB: 6AUD[1]). PIK3CA (PI3K*α*, PDB: 4JPS [2]), PIK3CB (PI3K*β*, PDB: 2Y3A [3]). The activation loop is shown in orange, with the k*α*12 helix shown in red (not a helix in PI3K*α*). The p85 regulatory subunits interacting with the motif in PI3K*α* and PI3K*β* are shown in blue, with the domains of the nSH2, iSH2, and cSH2 annotated on the structure. **(B)** Cartoon model shown in the same format as in Figure 1, highlighting the regulatory motif and its interaction with regulatory proteins.

**Figure S2 (relates to Fig 2).**
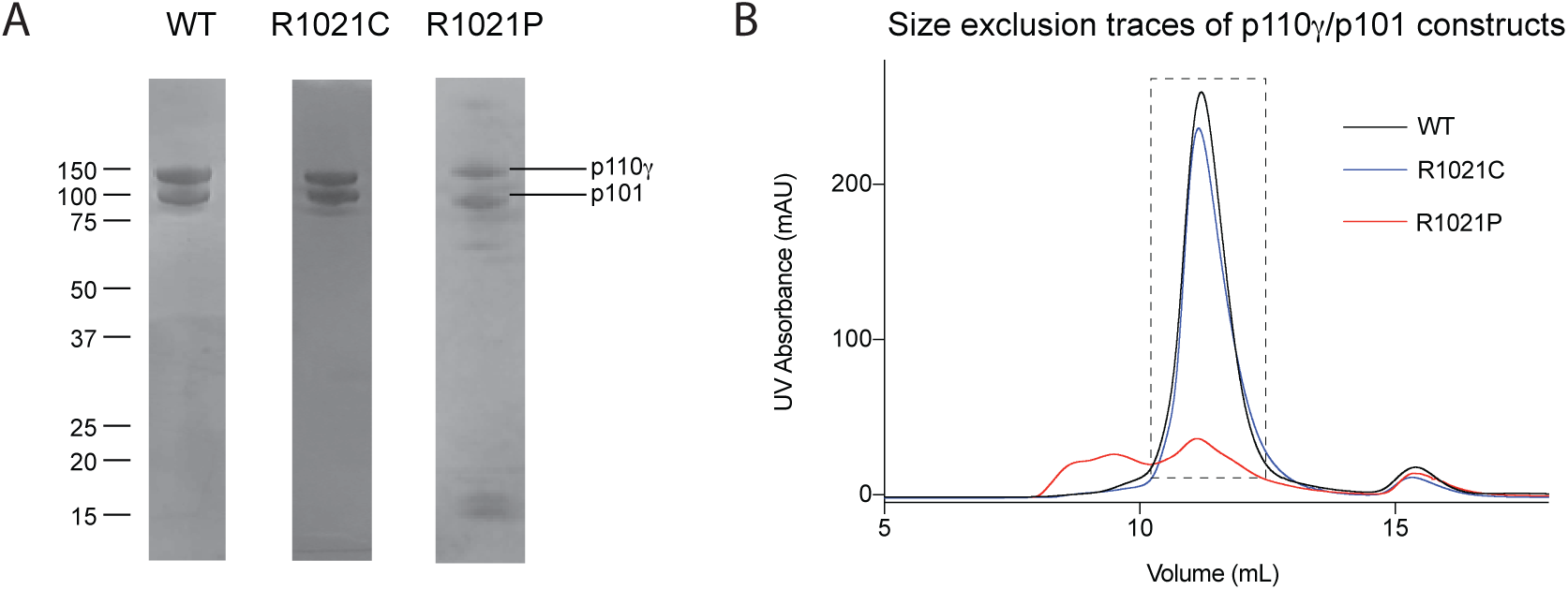
Purification of mutated p110γ / p101 complexes. **(A)** SDS-page analysis of the final complexes after size exclusion chromatography. The location of size markers are shown on the left. **(B)** Gel filtration elution of the wild type and mutant p110*γ* / p101 complexes on a Superdex™ 200 10/300 GL Increase column.

**Figure S3.**
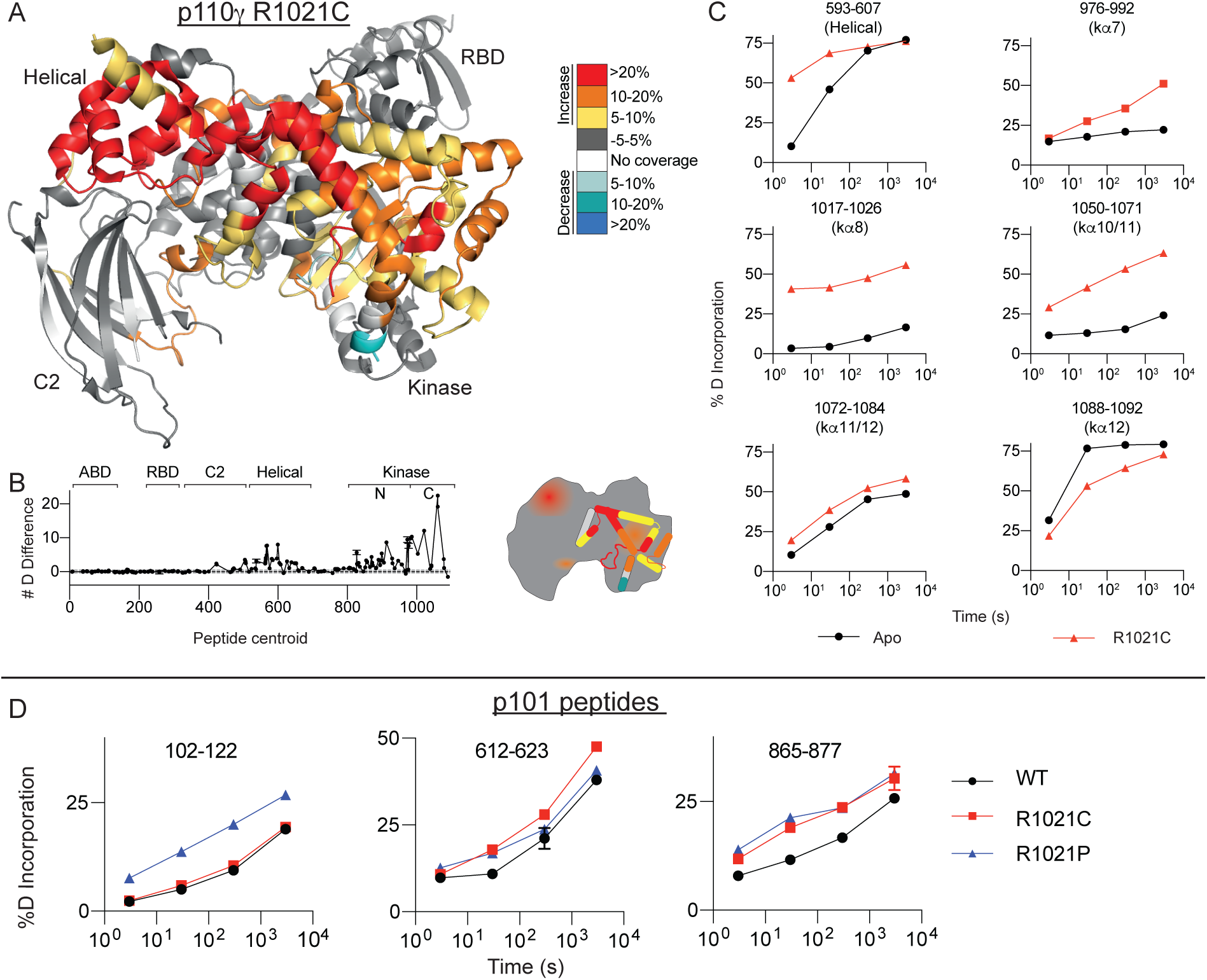
Differences in HDX for the R1021C mutation in free p110*γ*. **A.** Peptides showing significant deuterium exchange differences (>5 %, >0.4 kDa and p<0.01 in an unpaired two-tailed t-test) between p110*γ* wild-type and R1021C. Differences are colored on a model of p110*γ* (PDB: 6AUD). **B.** The number of deuteron difference for the R1021C mutant for all peptides analysed over the entire deuterium exchange time course for p110*γ*. **C.** Selected p110*γ* peptides that showed decreases and increases in exchange are shown. The full list of all peptides and their deuterium incorporation is shown in Supplementary Data 1. **D.** Selected p101 peptides that showed differences in exchange are shown. The full list of all peptides and their deuterium incorporation is shown in Supplementary Data 1.

**Figure S4.**
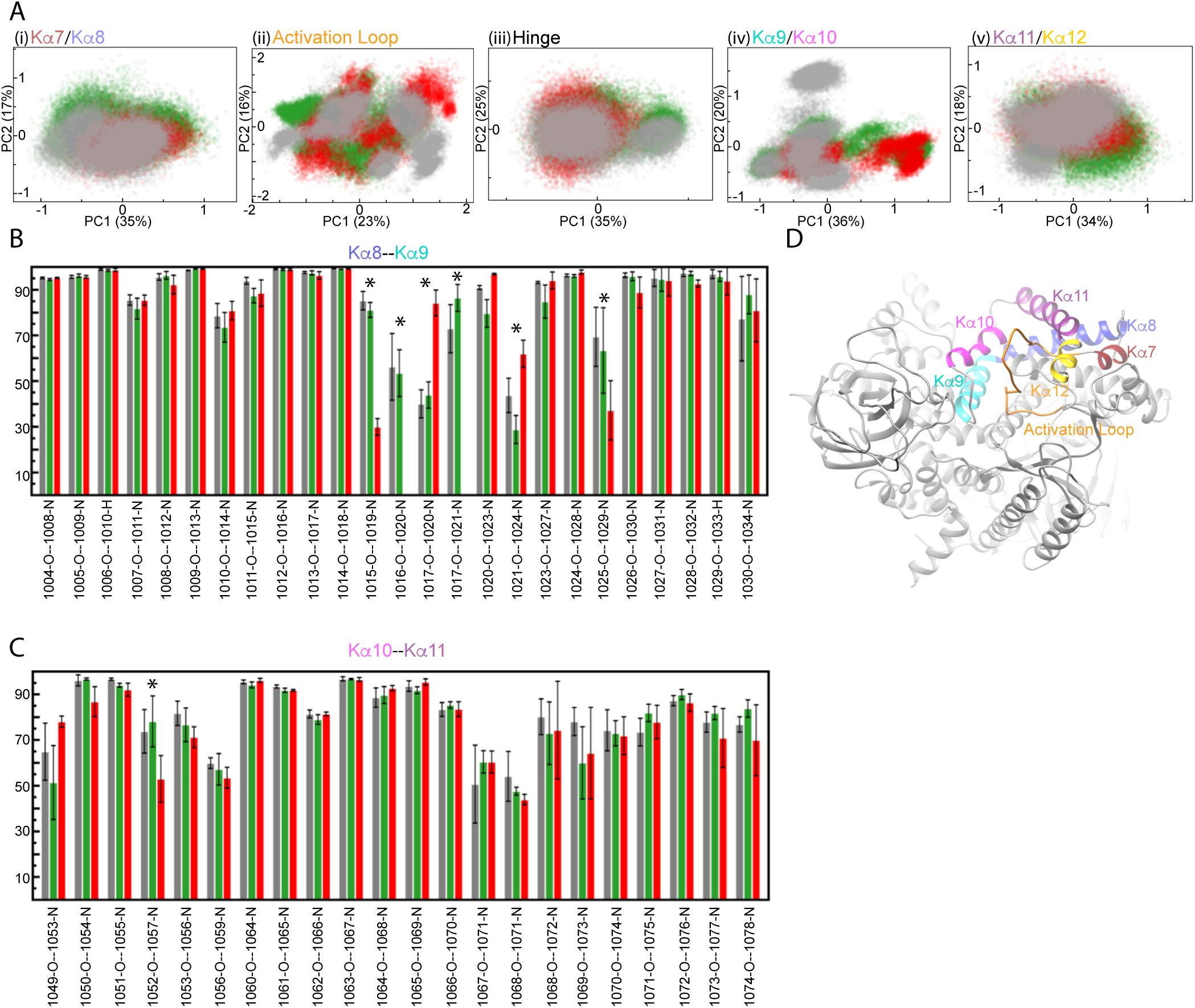
Differences between molecular dynamic simulations of WT, R1021C, and R1021P. **A.** Principal component analysis (PCA) plots showing PC1 vs. PC2 of K*α*7/8 (989-1023), Activation loop (962-988), hinge (879-887), k*α*9/10 (1024-1054) and k*α*11/12 (1057-1088) for WT (grey), R1021C (green) and R1021P (red) **B-C.** The mean and standard deviation of hydrogen bonding occupancies between k*α*8 and k*α*9 (B), k*α*10 and k*α*11 (C) across replicates for WT (grey), R1021C (green) and R1021P (red). Asterisks indicate significant differences between WT and mutants. **D.** Model of p110*γ* showing helices in the C-terminal regulatory motif and the activation loop.

**Figure S5.**
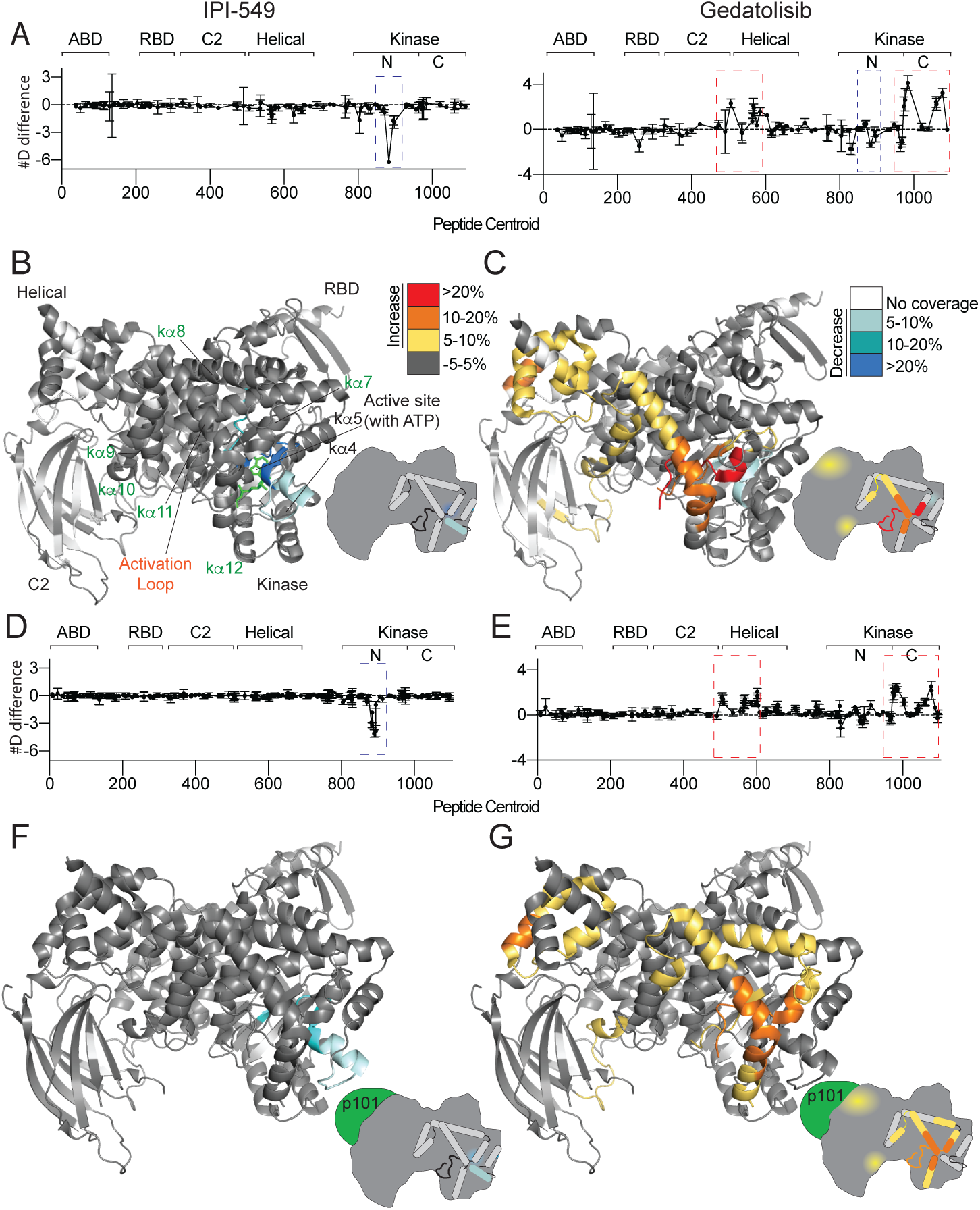
Differences in HDX for free p110*γ* and p110*γ*/p101 with selected inhibitors. **A.** Peptides showing significant deuterium exchange differences (>5 %, >0.4 kDa and p<0.01 in an unpaired two-tailed t-test) between p110*γ* wild-type and IPI-549 and Gedatolisib. Differences are colored on a model of p110*γ* (PDB: 6AUD). **B.** The number of deuteron difference for free p110*γ* with selected inhibitors for all peptides analysed over the entire deuterium exchange time course for p110*γ*. **C.** Peptides showing significant deuterium exchange differences (>5 %, >0.4 kDa and p<0.01 in an unpaired two-tailed t-test) between p110*γ*/p101 with IPI-549 and Gedatolisib. Differences are colored on a model of p110*γ* (PDB: 6AUD). **D.** The number of deuteron difference for p110*γ*/p101 with selected inhibitors for all peptides analysed over the entire deuterium exchange time course for p110*γ* and p101.

**Figure S6.**
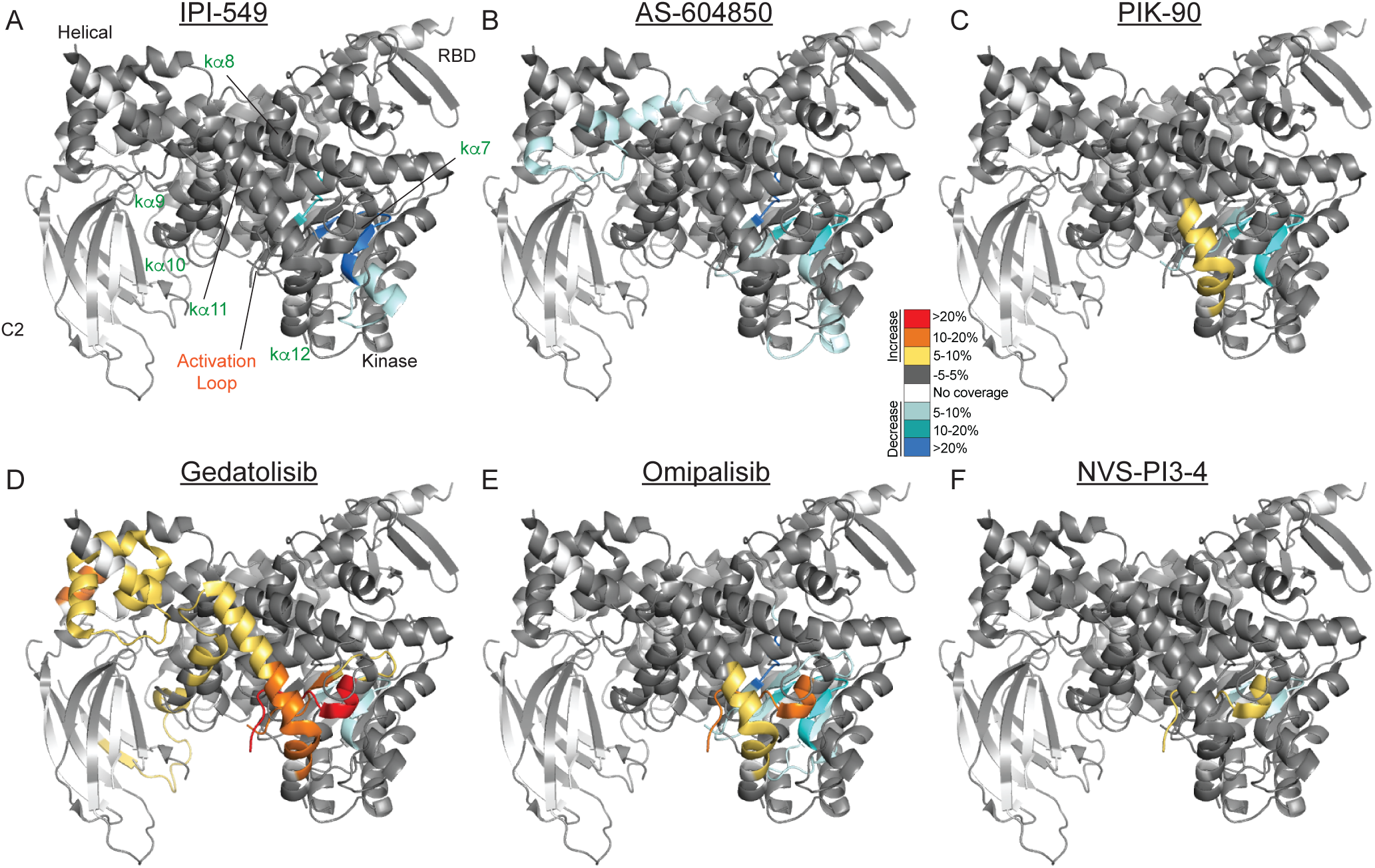
HDX-MS reveals that different classes of PI3K inhibitors lead to unique allosteric conformational changes. **A-F.** Peptides showing significant deuterium exchange differences (>5 %, >0.4 kDa and p<0.01 in an unpaired two-tailed t-test) between wild-type and six different inhibitors are colored on a model of p110*γ* (PDB: 6AUD). Differences in exchange are mapped according to the legend.

**Figure S7:**
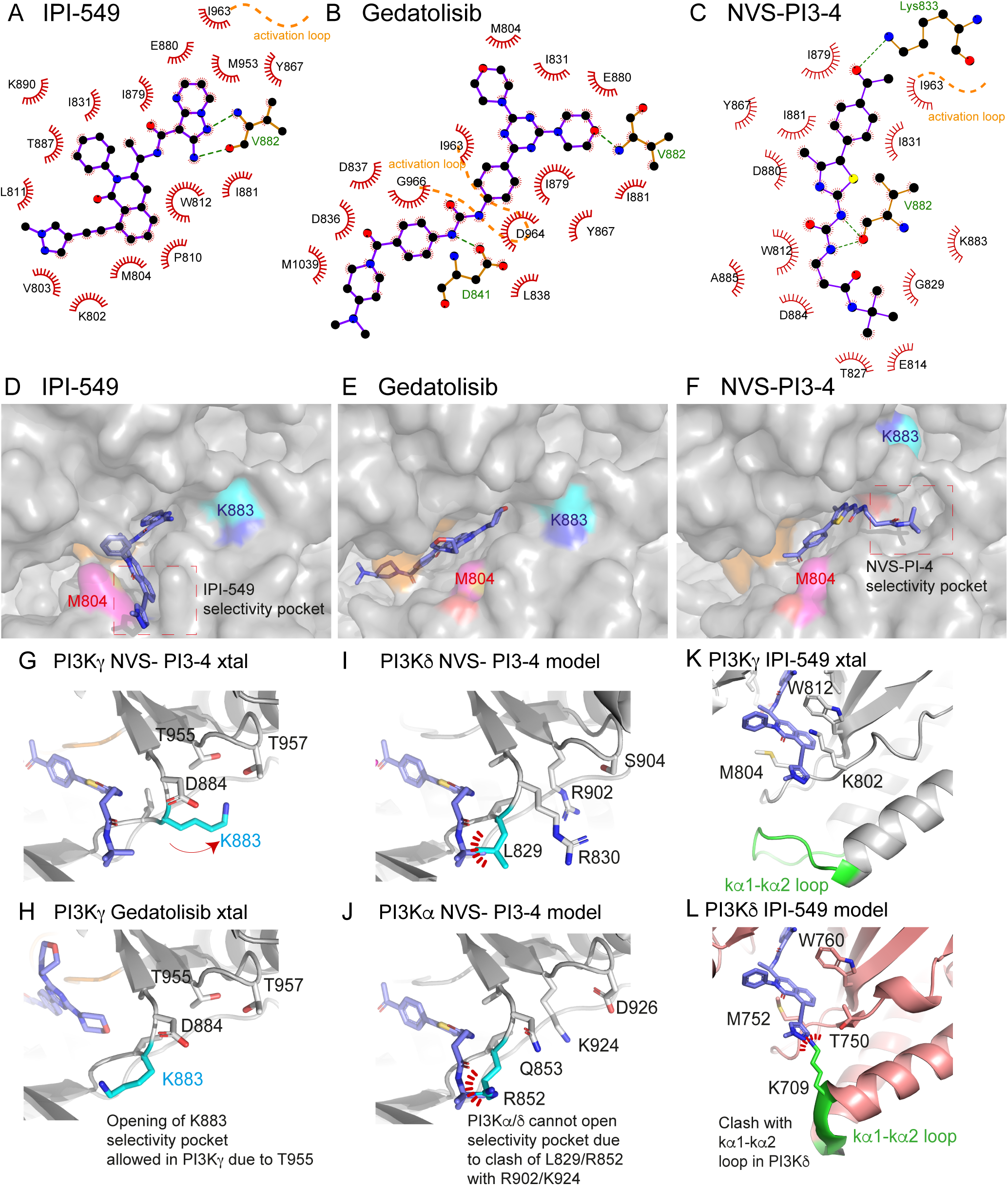
Structures of Gedatolisib and IPI-549 bound to p110γ. **A-C.** LigPlot+ [4] representations of p110*γ* bound to (**A**) IPI-549, (**B**) Gedatolisib, and (**C**) NVS-PI3-4. Hydrogen bonds are shown in green. All inhibitors form hydrogen bonds (green) with V882 in the hinge. The activation loop is shown as an orange dotted line. **D-E.** Comparison of Gedatolisib, IPI-549, and NVS-PI3-4 bound to p110*γ* with the activation loop and selectivity pockets highlighted. M804 and K883 that change conformation upon selectivity pocket opening are colored magenta and cyan, respectively. **G-J.** Molecular basis for NVS-PI3-4 for p110*γ* over p110*α*/*δ*. The structure of p110*γ* bound to NVS-PI3-4 (**G**) compared to p110*γ* bound to Gedatolisib (**H**), revealed a conformational change in K883 leading to opening of pocket accommodating the t-butyl motif. Comparing this to a model of p110*δ* (PDB: 5DXU) [5] (**I**) and p110*α* (PDB: 4JPS) [2] (**J**) with NVS-PI3-4 revealed that this pocket is unlikely to open with L829 in p110*δ* and R852 in p110*α* (corresponds to K883 in p110*γ*) unable to adopt this conformational change due to steric clashes / electrostatic repulsion with R902 in p110*δ* and K924 in p110*α* (corresponds to T955 in p110*γ*). **K-L.** Molecular basis for IPI-549 specificity for p110*γ* over p110*δ*. The structure of p110*γ* bound to IPI-549 (**H**) compared to a model of IPI-549 bound to p110*δ* (**I**), based on the structure of p110*δ* bound to the specificity pocket inhibitor Idelalisib (PDB: 4XE0) [6]. K802 and W812 in p110*γ* are labelled, along with the corresponding residues in p110*δ*. The k*α*1-k*α*2 loop is green, with potential clashes in p110*δ* with the methylpyrazole of IPI-549 highlighted.

**Fig. S8.**
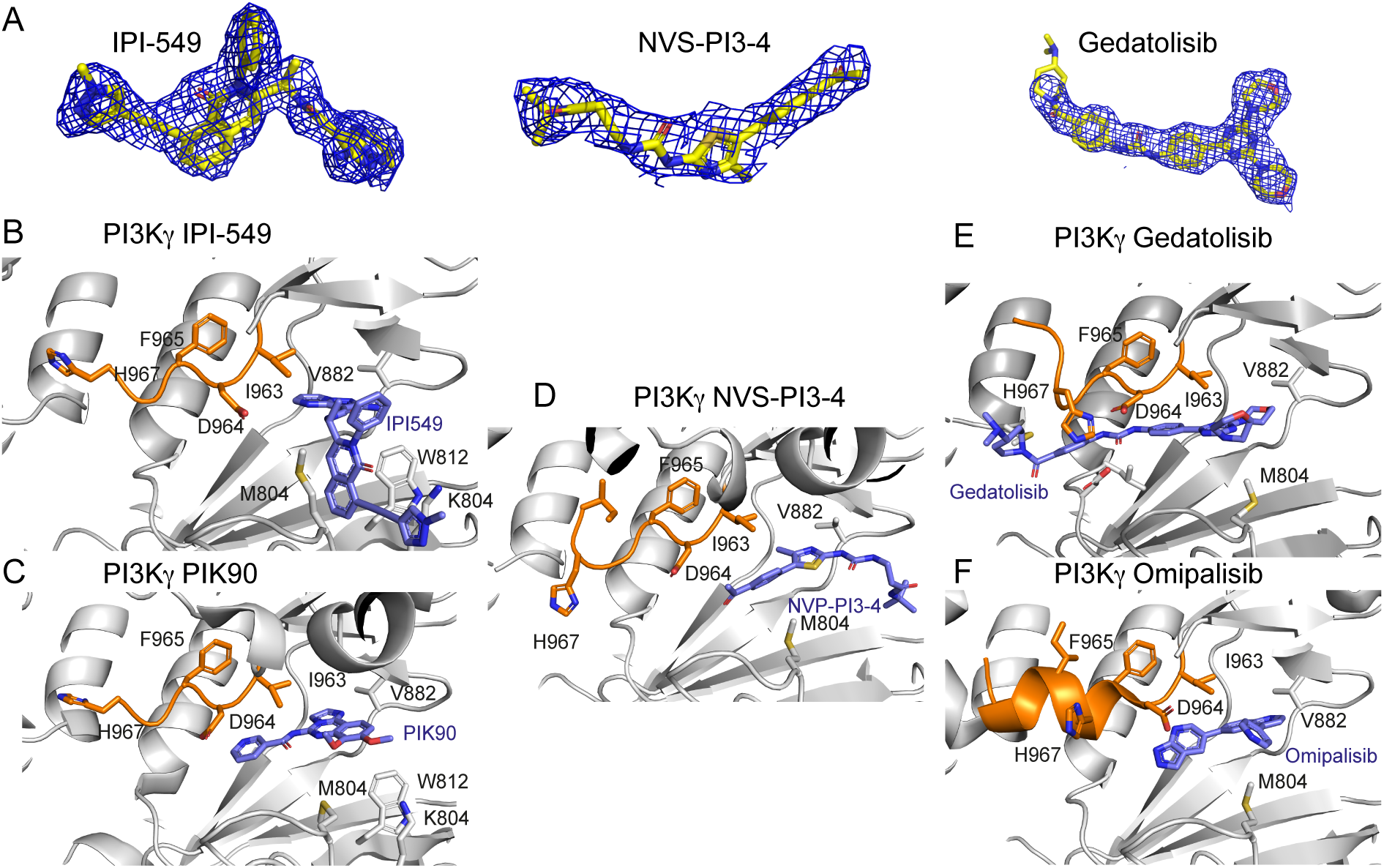
Binding of IPI-549, NVS-PI3-4, and Gedatolisib lead to different conformations of the activation loop of p110γ. **A.** The electron density from a feature enhanced map [7] around IPI-549, NVS-PI3-4, and Gedatolisib contoured at 1 sigma. **B-F.** Conformations of the activation loop of p110*γ* in the presence of annotated inhibitors. Structures of PIK90, and Omipalisib bound to p110*γ* were from PDB: 2CHX[8] and 3l54[9], respectively.

**Table S1.** Full HDX-MS experimental conditions and data analysis parameters from the guidelines of the IC-HDX-MS community [10].

| Data set | Apo p110γ<br>(mutants) | R1021C | p110γ / p101 | R1021C p110γ<br>p101 | R1021P p110γ<br>p101 |
| --- | --- | --- | --- | --- | --- |
| HDX reaction details | %D<sub>2</sub>O=87.9%<br>pH<sub>(read)</sub>=7.5<br>Temp=18°C | %D<sub>2</sub>O=87.9%<br>pH<sub>(read)</sub>=7.5<br>Temp=18°C | %D<sub>2</sub>O=62.0%<br>pH<sub>(read)</sub>=7.5<br>Temp=18°C | %D<sub>2</sub>O=62.0%<br>pH<sub>(read)</sub>=7.5<br>Temp=18°C | %D<sub>2</sub>O=62.0%<br>pH<sub>(read)</sub>=7.5<br>Temp=18°C |
| HDX time course (seconds) | 3, 30, 300, 3000 | 3, 30, 300, 3000 | 3, 30, 300, 3000 | 3, 30, 300, 3000 | 3, 30, 300, 3000 |
| HDX controls | N/A | N/A | N/A | N/A | N/A |
| Back-exchange | Corrected based on %D<sub>2</sub>O | Corrected based on %D<sub>2</sub>O | Corrected based on %D<sub>2</sub>O | Corrected based on %D<sub>2</sub>O | Corrected based on %D<sub>2</sub>O |
| Number of peptides | 204 | 202 | 153 | 153 | 152 |
| Sequence coverage | 92.7% | 92.5% | 92.7% | 92.7% | 91.7% |
| Average peptide /redundancy | Length=14.0<br>Redundancy= 2.4 | Length=14.0<br>Redundancy= 2.4 | Length=14.8<br>Redundancy= 2.0 | Length=14.8<br>Redundancy= 2.0 | Length=14.8<br>Redundancy= 2.0 |
| Replicates | 3 (2 3000s, 2 300s) | 3 (2 300s) | 3 | 3 | 3 |
| Repeatability | Average<br>StDev=0.5% | Average<br>StDev=0.5% | Average<br>StDev=0.6% | Average<br>StDev=0.6% | Average<br>StDev=0.6% |
| Significant differences in HDX | >5% and >0.4 Da and unpaired t-test $\leq 0.01$ | >5% and >0.4 Da and unpaired t-test $\leq 0.01$ | >5% and >0.4 Da and unpaired t-test $\leq 0.01$ | >5% and >0.4 Da and unpaired t-test $\leq 0.01$ | >5% and >0.4 Da and unpaired t-test $\leq 0.01$ |
| **Apo p110γ**<br>**(inhibitor)** | **+ IPI-549** | **+ AZg1/AZ** | **+ AS-605240** | **+ Gedatolisib** | **+ Omipalisib** |
| %D<sub>2</sub>O=75.5%<br>pH<sub>(read)</sub>=7.5<br>Temp=18°C | %D<sub>2</sub>O=75.5%<br>pH<sub>(read)</sub>=7.5<br>Temp=18°C | %D<sub>2</sub>O=75.5%<br>pH<sub>(read)</sub>=7.5<br>Temp=18°C | %D<sub>2</sub>O=75.5%<br>pH<sub>(read)</sub>=7.5<br>Temp=18°C | %D<sub>2</sub>O=75.5%<br>pH<sub>(read)</sub>=7.5<br>Temp=18°C | %D<sub>2</sub>O=75.5%<br>pH<sub>(read)</sub>=7.5<br>Temp=18°C |
| 3, 30, 300, 3000 | 3, 30, 300, 3000 | 3, 30, 300, 3000 | 3, 30, 300, 3000 | 3, 30, 300, 3000 | 3, 30, 300, 3000 |
| N/A | N/A | N/A | N/A | N/A | N/A |
| Corrected based on %D<sub>2</sub>O | Corrected based on %D<sub>2</sub>O | Corrected based on %D<sub>2</sub>O | Corrected based on %D<sub>2</sub>O | Corrected based on %D<sub>2</sub>O | Corrected based on %D<sub>2</sub>O |
| 180 | 180 | 180 | 180 | 180 | 180 |
| 88.6% | 88.6% | 88.6% | 88.6% | 88.6% | 88.6% |
| Length= 13.4<br>Redundancy= 2.2 | Length= 13.4<br>Redundancy= 2.2 | Length= 13.4<br>Redundancy= 2.2 | Length= 13.4<br>Redundancy= 2.2 | Length= 13.4<br>Redundancy= 2.2 | Length= 13.4<br>Redundancy= 2.2 |
| 3 | 3 | 3 | 3 | 3 | 3 |
| Average<br>StDev=0.9% | Average<br>StDev=0.9% | Average<br>StDev=0.9% | Average<br>StDev=0.9% | Average<br>StDev=0.9% | Average<br>StDev=0.9% |
| >5% and >0.4 Da and unpaired t-test $\leq 0.01$ | >5% and >0.4 Da and unpaired t-test $\leq 0.01$ | >5% and >0.4 Da and unpaired t-test $\leq 0.01$ | >5% and >0.4 Da and unpaired t-test $\leq 0.01$ | >5% and >0.4 Da and unpaired t-test $\leq 0.01$ | >5% and >0.4 Da and unpaired t-test $\leq 0.01$ |
| **+ RD-HBC 520** | **+ PIK-90** | **Apo p110γ/p101**<br>**(p101+inhibitor)** | **+ IPI-549 (p101)** | **+ Gedatolisib (p101)** |  |
| %D<sub>2</sub>O=75.5%<br>pH<sub>(read)</sub>=7.5<br>Temp=18°C | %D<sub>2</sub>O=75.5%<br>pH<sub>(read)</sub>=7.5<br>Temp=18°C | %D<sub>2</sub>O=75.5%<br>pH<sub>(read)</sub>=7.5<br>Temp=18°C | %D<sub>2</sub>O=75.5%<br>pH<sub>(read)</sub>=7.5<br>Temp=18°C | %D<sub>2</sub>O=75.5%<br>pH<sub>(read)</sub>=7.5<br>Temp=18°C |  |
| 3, 30, 300, 3000 | 3, 30, 300, 3000 | 3, 30, 300, 3000 | 3, 30, 300, 3000 | 3, 30, 300, 3000 |  |
| N/A | N/A | N/A | N/A | N/A |  |
| Corrected based on %D<sub>2</sub>O | Corrected based on %D<sub>2</sub>O | Corrected based on %D<sub>2</sub>O | Corrected based on %D<sub>2</sub>O | Corrected based on %D<sub>2</sub>O |  |
| 180 | 180 | 228 | 228 | 228 |  |
| 88.6% | 88.6% | 96.3% | 96.3% | 96.3% |  |
| Length= 13.4<br>Redundancy= 2.2 | Length= 13.4<br>Redundancy= 2.2 | Length= 14.1<br>Redundancy= 2.9 | Length= 14.1<br>Redundancy= 2.9 | Length= 13.4<br>Redundancy= 2.2 |  |
| 3 | 3 | 3 | 3 | 3 |  |
| Average<br>StDev=0.9% | Average<br>StDev=0.9% | Average<br>StDev=0.9% | Average<br>StDev=0.9% | Average<br>StDev=0.9% |  |
| >5% and >0.4 Da and unpaired t-test $\leq 0.01$ | >5% and >0.4 Da and unpaired t-test $\leq 0.01$ | >5% and >0.4 Da and unpaired t-test $\leq 0.01$ | >5% and >0.4 Da and unpaired t-test $\leq 0.01$ | >5% and >0.4 Da and unpaired t-test $\leq 0.01$ | |

**Table S2.**
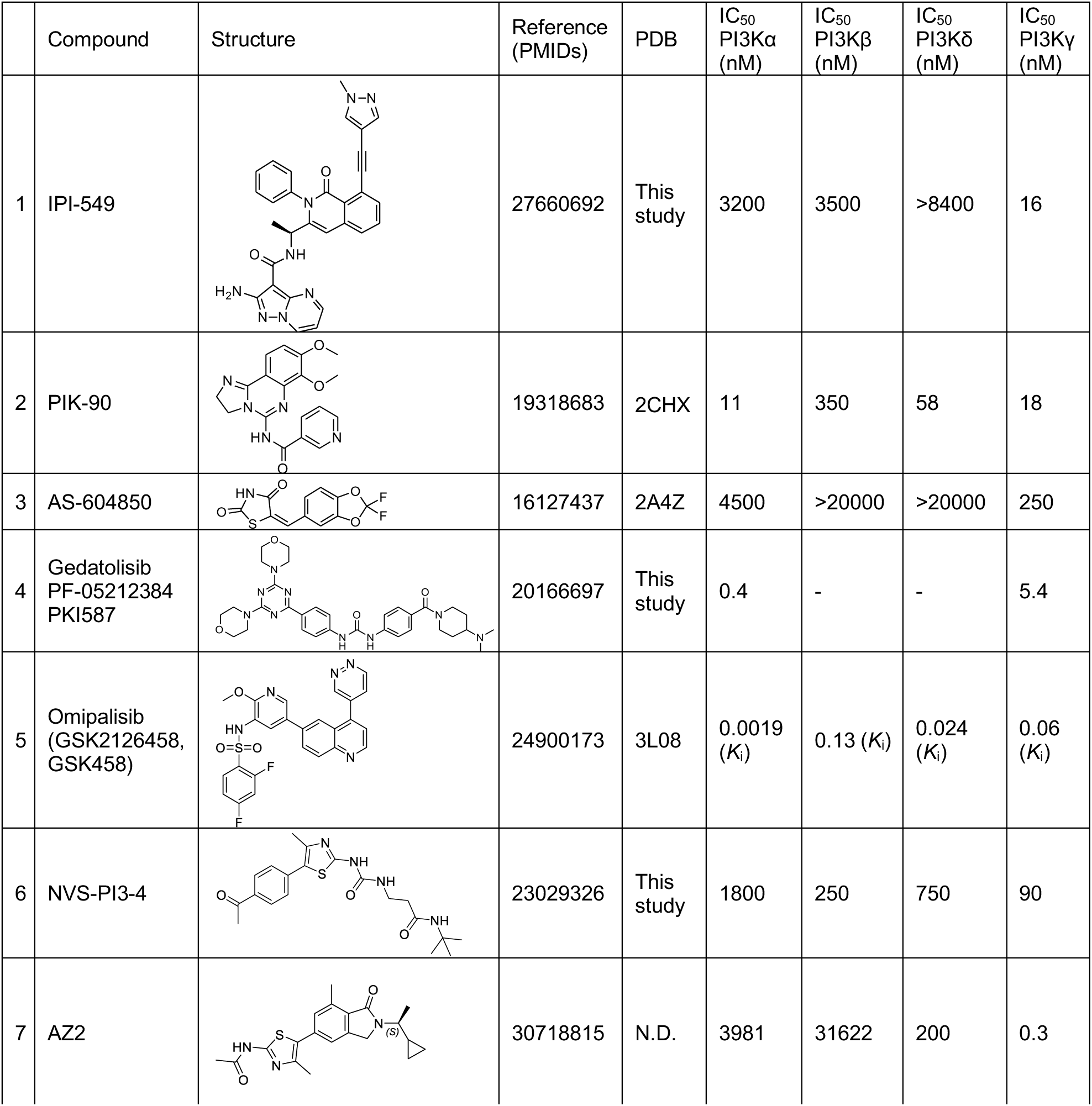
List of all PI3K inhibitors analysed in this manuscript. IC50s for class IA and IB are listed from the reference attached. N.D. is not determined.

| | Compound | Structure | Reference (PMIDs) | PDB | IC<sub>50</sub> PI3Kα (nM) | IC<sub>50</sub> PI3Kβ (nM) | IC<sub>50</sub> PI3Kδ (nM) | IC<sub>50</sub> PI3Kγ (nM) |
| --- | --- | --- | --- | --- | --- | --- | --- | --- |
| 1 | IPI-549 |  | 27660692 | This study | 3200 | 3500 | >8400 | 16 |
| 2 | PIK-90 |  | 19318683 | 2CHX | 11 | 350 | 58 | 18 |
| 3 | AS-604850 |  | 16127437 | 2A4Z | 4500 | >20000 | >20000 | 250 |
| 4 | Gedatolisib<br>PF-05212384<br>PKI587 |  | 20166697 | This study | 0.4 | - | - | 5.4 |
| 5 | Omipalisib<br>(GSK2126458,<br>GSK458) |  | 24900173 | 3L08 | 0.0019<br>(K<sub>i</sub>) | 0.13 (K<sub>i</sub>) | 0.024<br>(K<sub>i</sub>) | 0.06<br>(K<sub>i</sub>) |
| 6 | NVS-PI3-4 |  | 23029326 | This study | 1800 | 250 | 750 | 90 |
| 7 | AZ2 |  | 30718815 | N.D. | 3981 | 31622 | 200 | 0.3 |

**Table S3.** X-ray Data collection and refinement statistics

| | PI3Kγ<br>IPI549 | PI3Kγ<br>Gedatolisib | PI3Kγ<br>NVS-PI3-4 |
| --- | --- | --- | --- |
| **Data collection** |  |  |  |
| Wavelength | 0.97949 | 0.97949 | 0.97949 |
| Space group | C121 | C121 | C121 |
| Cell dimensions |  |  |  |
| $a, b, c$ (Å) | 144.3, 67.9, 106.4 | 143.5, 67.6, 106.3 | 143.6 67.6 106.8 |
| $\alpha, \beta, \gamma$ (°) | 90 94.5 90 | 90, 95.4, 90 | 90 95.4 90 |
| Resolution (Å) | 44.4 - 2.65 (2.74 - 2.65)* | 40.72-2.55 (2.64-2.55) | 40.93 - 3.15 (3.26 - 3.15) |
| $R_{\text{merge}}$ | 0.125 (1.919) | 0.061 (1.349) | 0.119 (1.118) |
| $I / \sigma I$ | 7.1 (0.69) | 11.91 (0.87) | 7.92 (0.84) |
| CC1/2 | 0.992 (0.407) | 0.999 (0.385) | 0.994 (0.425) |
| Completeness (%) | 98.9 (98.23) | 99.41 (99.40) | 98.08 (99.04) |
| Redundancy | 3.3 (3.4) | 3.3 (3.4) | 3.0 (3.0) |
| **Refinement** |  |  |  |
| Resolution (Å) | 44.4 - 2.65 (2.74 - 2.65) | 40.72-2.55 (2.64-2.55) | 40.93 - 3.15 (3.26 - 3.15) |
| No. unique reflections | 29722 (2941) | 33183 (3303) | 17573 (1761) |
| $R_{\text{work}} / R_{\text{free}}$ | 22.7/26.8 | 20.9/25.3 | 22.9/27.4 |
| No. atoms |  |  |  |
| Protein | 6752 | 6612 | 6506 |
| Ligand/ion | 40 | 45 | 28 |
| Water | 0 | 9 | 0 |
| $B$-factors | | | |
| Protein | 100.4 | 88.9 | 108.2 |
| Ligand/ion | 88.3 | 78.7 | 117.2 |
| Water |  | 65.5 |  |
| Ramachandran favored | 94.47 | 95.21 | 96.51 |
| Ramachandran outliers | 0.61 | 0.0 | 0.13 |
| Rotamer outliers | 0.53 | 0.41 | 0.0 |
| R.m.s. deviations |  |  |  |
| Bond lengths (Å) | 0.003 | 0.003 | 0.004 |
| Bond angles (°) | 0.53 | 0.59 | 0.56 |

\*Values in parentheses are for highest-resolution shell.
Number of crystals used for structure=1

**Source data figure legend. Summary of all HDX-MS peptide data (see attached excel source data file).** The charge state (Z), residue start (S), residue end number (E), and retention time (RT) are displayed for every peptide. Data listed is the mean of 3 independent experiments, with SDs presented. Time points are labelled, and the relative level of HDX is coloured according to the legend.

## References

1. Bilanges B, Posor Y, Vanhaesebroeck B. PI3K isoforms in cell signalling and vesicle trafficking. Nat Rev Mol Cell Biol. Nature Publishing Group; 2019 May 20;10:6742.

2. Madsen RR, Vanhaesebroeck B. Cracking the context-specific PI3K signaling code. Sci Signal. American Association for the Advancement of Science; 2020 Jan 7;13(613):eaay2940.

3. Suire S, Coadwell J, Ferguson GJ, Davidson K, Hawkins P, Stephens L. p84, a new Gbetagamma-activated regulatory subunit of the type IB phosphoinositide 3- kinase p110gamma. Curr Biol [Internet]. 2005 Mar 29;15(6):566–70. Available from: http://www.ncbi.nlm.nih.gov/entrez/query.fcgi?cmd=Retrieve&db=PubMed&dopt=Citation&list_uids=15797027

4. Stephens LR, Eguinoa A, Erdjument-Bromage H, Lui M, Cooke F, Coadwell J, Smrcka AS, Thelen M, Cadwallader K, Tempst P, Hawkins PT. The G beta gamma sensitivity of a PI3K is dependent upon a tightly associated adaptor, p101. Cell. 1997 Apr 4;89(1):105–14.

5. Bohnacker T, Marone R, Collmann E, Calvez R, Hirsch E, Wymann M. PI3Kgamma adaptor subunits define coupling to degranulation and cell motility by distinct PtdIns(3,4,5)P3 pools in mast cells. Sci Signal. 2009 Jan;2(74):ra27.

6. Goncalves MD, Cantley LC. Phosphatidylinositol 3-Kinase, Growth Disorders, and Cancer. Vol. 379, The New England journal of medicine. 2018. pp. 2052–62.

7. Fruman DA, Chiu H, Hopkins BD, Bagrodia S, Cantley LC, Abraham RT. The PI3K Pathway in Human Disease. Cell. 2017 Aug 10;170(4):605–35.

8. Burke JE. Structural Basis for Regulation of Phosphoinositide Kinases and Their Involvement in Human Disease. Mol Cell. 2018 Sep 6;71(5):653–73.

9. Camps M, Rückle T, Ji H, Ardissone V, Rintelen F, Shaw J, Ferrandi C, Chabert C, Gillieron C, Françon B, Martin T, Gretener D, Perrin D, Leroy D, Vitte P-A, Hirsch E, Wymann MP, Cirillo R, Schwarz MK, Rommel C. Blockade of PI3Kgamma suppresses joint inflammation and damage in mouse models of rheumatoid arthritis. Nat Med. 2005 Sep;11(9):936–43.

10. Patrucco E, Notte A, Barberis L, Selvetella G, Maffei A, Brancaccio M, Marengo S, Russo G, Azzolino O, Rybalkin SD, Silengo L, Altruda F, Wetzker R, Wymann MP, Lembo G, Hirsch E. PI3Kgamma modulates the cardiac response to chronic pressure overload by distinct kinase-dependent and -independent effects. Cell. 2004 Aug 6;118(3):375–87.

11. Hirsch E, Katanaev VL, Garlanda C, Azzolino O, Pirola L, Silengo L, Sozzani S, Mantovani A, Altruda F, Wymann MP. Central role for G protein-coupled phosphoinositide 3-kinase gamma in inflammation. Science. 2000 Feb 11;287(5455):1049–53.

12. Li Z, Jiang H, Xie W, Zhang Z, Smrcka AV, Wu D. Roles of PLC-beta2 and -beta3 and PI3Kgamma in chemoattractant-mediated signal transduction. Science. American Association for the Advancement of Science; 2000 Feb 11;287(5455):1046–9.

13. Sasaki T, Irie-Sasaki J, Jones RG, Oliveira-dos-Santos AJ, Stanford WL, Bolon B, Wakeham A, Itie A, Bouchard D, Kozieradzki I, Joza N, Mak TW, Ohashi PS, Suzuki A, Penninger JM. Function of PI3Kgamma in thymocyte development, T cell activation, and neutrophil migration. Science. American Association for the Advancement of Science; 2000 Feb 11;287(5455):1040–6.

14. Laffargue M, Calvez R, Finan P, Trifilieff A, Barbier M, Altruda F, Hirsch E, Wymann MP. Phosphoinositide 3-kinase gamma is an essential amplifier of mast cell function. Immunity. 2002 Mar;16(3):441–51.

15. Collmann E, Bohnacker T, Marone R, Dawson J, Rehberg M, Stringer R, Krombach F, Burkhart C, Hirsch E, Hollingworth GJ, Thomas M, Wymann MP. Transient targeting of phosphoinositide 3-kinase acts as a roadblock in mast cells’ route to allergy. J Allergy Clin Immunol. 2013 Oct;132(4):959–68.

16. Stoyanov B, Volinia S, Hanck T, Rubio I, Loubtchenkov M, Malek D, Stoyanova S, Vanhaesebroeck B, Dhand R, Nurnberg B. Cloning and characterization of a G protein-activated human phosphoinositide-3 kinase. Science. 1995 Aug 4;269(5224):690–3.

17. Schmid MC, Avraamides CJ, Dippold HC, Franco I, Foubert P, Ellies LG, Acevedo LM, Manglicmot JRE, Song X, Wrasidlo W, Blair SL, Ginsberg MH, Cheresh DA, Hirsch E, Field SJ, Varner JA. Receptor tyrosine kinases and TLR/IL1Rs unexpectedly activate myeloid cell PI3kγ, a single convergent point promoting tumor inflammation and progression. Cancer Cell. 2011 Jun 14;19(6):715–27.

18. Luo L, Wall AA, Tong SJ, Hung Y, Xiao Z, Tarique AA, Sly PD, Fantino E, Marzolo M-P, Stow JL. TLR Crosstalk Activates LRP1 to Recruit Rab8a and PI3Kγ for Suppression of Inflammatory Responses. Cell Rep. 2018 Sep 11;24(11):3033–44.

19. Luo L, Wall AA, Yeo JC, Condon ND, Norwood SJ, Schoenwaelder S, Chen KW, Jackson S, Jenkins BJ, Hartland EL, Schroder K, Collins BM, Sweet MJ, Stow JL. Rab8a interacts directly with PI3Kγ to modulate TLR4-driven PI3K and mTOR signalling. Nat Commun. 2014;5:4407.

20. Stephens LR, Eguinoa A, Erdjument-Bromage H, Lui M, Cooke F, Coadwell J, Smrcka AS, Thelen M, Cadwallader K, Tempst P, Hawkins PT. The Gbg sensitivity of a PI3K is dependent upon a tightly associated adaptor, p101. Cell. 1997 Jan;89:105–14.

21. Vadas O, Dbouk HA, Shymanets A, Perisic O, Burke JE, Abi Saab WF, Khalil BD, Harteneck C, Bresnick AR, Nürnberg B, Backer JM, Williams RL. Molecular determinants of PI3Kγ-mediated activation downstream of G-protein-coupled receptors (GPCRs). Proc Natl Acad Sci USA. National Acad Sciences; 2013 Nov 19;110(47):18862–7.

22. Kurig B, Shymanets A, Bohnacker T, Prajwal, Brock C, Ahmadian MR, Schaefer M, Gohla A, Harteneck C, Wymann MP, Jeanclos E, Nürnberg B. Ras is an indispensable coregulator of the class IB phosphoinositide 3-kinase p87/p110gamma. Proc Natl Acad Sci USA [Internet]. 2009 Nov 20;106(48):20312–7. Available from: http://www.ncbi.nlm.nih.gov/entrez/query.fcgi?cmd=Retrieve&db=PubMed&dopt=Citation&list_uids=19906996

23. Burke JE, Williams RL. Synergy in activating class I PI3Ks. Trends in Biochemical Sciences. 2015 Feb;40(2):88–100.

24. Barber D, Bartolome A, Hernandez C, Flores J, Redondo C, Fernandez-Arias C, Camps M, Ruckle T, Schwarz M, Rodriguez S, Martinez AC, Balomenos D, Rommel C, Carrera A. PI3Kgamma inhibition blocks glomerulonephritis and extends lifespan in a mouse model of systemic lupus. Nat Med [Internet]. 2005 Aug 20;11(9):933–5. Available from: http://www.ncbi.nlm.nih.gov/entrez/query.fcgi?cmd=Retrieve&db=PubMed&dopt=Citation&list_uids=16127435

25. Thomas M, Edwards MJ, Sawicka E, Duggan N, Hirsch E, Wymann MP, Owen C, Trifilieff A, Walker C, Westwick J, Finan P. Essential role of phosphoinositide 3- kinase gamma in eosinophil chemotaxis within acute pulmonary inflammation. Immunology. John Wiley & Sons, Ltd; 2009 Mar;126(3):413–22.

26. Campa CC, Silva RL, Margaria JP, Pirali T, Mattos MS, Kraemer LR, Reis DC, Grosa G, Copperi F, Dalmarco EM, Lima-Júnior RCP, Aprile S, Sala V, Dal Bello F, Prado DS, Alves-Filho JC, Medana C, Cassali GD, Tron GC, Teixeira MM, Ciraolo E, Russo RC, Hirsch E. Inhalation of the prodrug PI3K inhibitor CL27c improves lung function in asthma and fibrosis. Nat Commun. Nature Publishing Group; 2018 Dec 12;9(1):5232–16.

27. Breasson L, Becattini B, Sardi C, Molinaro A, Zani F, Marone R, Botindari F, Bousquenaud M, Ruegg C, Wymann MP, Solinas G. PI3Kγ activity in leukocytes promotes adipose tissue inflammation and early-onset insulin resistance during obesity. Sci Signal. American Association for the Advancement of Science; 2017 Jul 18;10(488):eaaf2969.

28. Kaneda MM, Cappello P, Nguyen AV, Ralainirina N, Hardamon CR, Foubert P, Schmid MC, Sun P, Mose E, Bouvet M, Lowy AM, Valasek MA, Sasik R, Novelli F, Hirsch E, Varner JA. Macrophage PI3Kγ Drives Pancreatic Ductal Adenocarcinoma Progression. Cancer Discov. American Association for Cancer Research; 2016 Aug;6(8):870–85.

29. De Henau O, Rausch M, Winkler D, Campesato LF, Liu C, Cymerman DH, Budhu S, Ghosh A, Pink M, Tchaicha J, Douglas M, Tibbitts T, Sharma S, Proctor J, Kosmider N, White K, Stern H, Soglia J, Adams J, Palombella VJ, McGovern K, Kutok JL, Wolchok JD, Merghoub T. Overcoming resistance to checkpoint blockade therapy by targeting PI3Kγ in myeloid cells. Nature. 2016 Nov 9.

30. Kaneda MM, Messer KS, Ralainirina N, Li H, Leem CJ, Gorjestani S, Woo G, Nguyen AV, Figueiredo CC, Foubert P, Schmid MC, Pink M, Winkler DG, Rausch M, Palombella VJ, Kutok J, McGovern K, Frazer KA, Wu X, Karin M, Sasik R, Cohen EEW, Varner JA. PI3Kγ is a molecular switch that controls immune suppression. Nature. 2016 Nov 17;539(7629):437–42.

31. Walker EH, Perisic O, Ried C, Stephens L, Williams RL. Structural insights into phosphoinositide 3-kinase catalysis and signalling. Nature. 1999 Nov 18;402(6759):313–20.

32. Deladeriere A, Gambardella L, Pan D, Anderson KE, Hawkins PT, Stephens LR. The regulatory subunits of PI3Kγ control distinct neutrophil responses. Sci Signal. 2015;8(360):ra8.

33. Pacold ME, Suire S, Perisic O, Lara-Gonzalez S, Davis CT, Walker EH, Hawkins PT, Stephens L, Eccleston JF, Williams RL. Crystal structure and functional analysis of Ras binding to its effector phosphoinositide 3-kinase gamma. Cell [Internet]. 2000 Dec 8;103(6):931–43. Available from: http://www.ncbi.nlm.nih.gov/entrez/query.fcgi?cmd=Retrieve&db=PubMed&dopt=Citation&list_uids=11136978

34. Gangadhara G, Dahl G, Bohnacker T, Rae R, Gunnarsson J, Blaho S, Öster L, Lindmark H, Karabelas K, Pemberton N, Tyrchan C, Mogemark M, Wymann MP, Williams RL, Perry MWD, Papavoine T, Petersen J. A class of highly selective inhibitors bind to an active state of PI3Kγ. Nature Chemical Biology. Nature Publishing Group; 2019 Apr;15(4):348–57.

35. Vadas O, Burke JE, Zhang X, Berndt A, Williams RL. Structural basis for activation and inhibition of class I phosphoinositide 3-kinases. Sci Signal [Internet]. 2011 Oct 18;4(195):1–13. Available from: http://www.ncbi.nlm.nih.gov/entrez/query.fcgi?cmd=Retrieve&db=PubMed&dopt=Citation&list_uids=22009150

36. Kang S, Denley A, Vanhaesebroeck B, Vogt PK. Oncogenic transformation induced by the p110beta, -gamma, and -delta isoforms of class I phosphoinositide 3-kinase. Proc Natl Acad Sci USA. 2006 Jan 31;103(5):1289–94.

37. Samuels Y, Wang Z, Bardelli A, Silliman N, Ptak J, Szabo S, Yan H, Gazdar A, Powell S, Riggins G, Willson J, Markowitz S, Kinzler K, Vogelstein B, Velculescu V. High frequency of mutations of the PIK3CA gene in human cancers. Science. 2004 May 23;304(5670):554.

38. Vasan N, Razavi P, Johnson JL, Shao H, Shah H, Antoine A, Ladewig E, Gorelick A, Lin T-Y, Toska E, Xu G, Kazmi A, Chang MT, Taylor BS, Dickler MN, Jhaveri K, Chandarlapaty S, Rabadan R, Reznik E, Smith ML, Sebra R, Schimmoller F, Wilson TR, Friedman LS, Cantley LC, Scaltriti M, Baselga J. Double PIK3CA mutations in cis increase oncogenicity and sensitivity to PI3Kα inhibitors. Science. American Association for the Advancement of Science; 2019 Nov 8;366(6466):714–23.

39. Lindhurst MJ, Parker VER, Payne F, Sapp JC, Rudge S, Harris J, Witkowski AM, Zhang Q, Groeneveld MP, Scott CE, Daly A, Huson SM, Tosi LL, Cunningham ML, Darling TN, Geer J, Gucev Z, Sutton VR, Tziotzios C, Dixon AK, Helliwell T, O’Rahilly S, Savage DB, Wakelam MJO, Barroso I, Biesecker LG, Semple RK. Mosaic overgrowth with fibroadipose hyperplasia is caused by somatic activating mutations in PIK3CA. Nat Genet. 2012 Aug;44(8):928–33.

40. Dornan GL, Siempelkamp BD, Jenkins ML, Vadas O, Lucas CL, Burke JE. Conformational disruption of PI3Kδ regulation by immunodeficiency mutations in PIK3CD and PIK3R1. Proc Natl Acad Sci USA. 2017 Feb 21;114(8):1982–7.

41. Lucas CL, Chandra A, Nejentsev S, Condliffe AM, Okkenhaug K. PI3Kδ and primary immunodeficiencies. Nat Rev Immunol. 2016 Nov;16(11):702–14.

42. Angulo I, Vadas O, Garçon F, Banham-Hall E, Plagnol V, Leahy TR, Baxendale H, Coulter T, Curtis J, Wu C, Blake-Palmer K, Perisic O, Smyth D, Maes M, Fiddler C, Juss J, Cilliers D, Markelj G, Chandra A, Farmer G, Kielkowska A, Clark J, Kracker S, Debré M, Picard C, Pellier I, Jabado N, Morris JA, Barcenas-Morales G, Fischer A, Stephens L, Hawkins P, Barrett JC, Abinun M, Clatworthy M, Durandy A, Doffinger R, Chilvers ER, Cant AJ, Kumararatne D, Okkenhaug K, Williams RL, Condliffe A, Nejentsev S. Phosphoinositide 3-kinase δ gene mutation predisposes to respiratory infection and airway damage. Science. American Association for the Advancement of Science; 2013 Nov 15;342(6160):866–71.

43. Lowery MA, Bradley M, Chou JF, Capanu M, Gerst S, Harding JJ, Dika IE, Berger M, Zehir A, Ptashkin R, Wong P, Rasalan-Ho T, Yu KH, Cercek A, Morgono E, Salehi E, Valentino E, Hollywood E, O’Reilly EM, Abou-Alfa GK. Binimetinib plus Gemcitabine and Cisplatin Phase I/II Trial in Patients with Advanced Biliary Cancers. Clin Cancer Res. American Association for Cancer Research; 2019 Feb 1;25(3):937–45.

44. AACR Project GENIE Consortium. AACR Project GENIE: Powering Precision Medicine through an International Consortium. Cancer Discov. American Association for Cancer Research; 2017 Aug;7(8):818–31.

45. Tate JG, Bamford S, Jubb HC, Sondka Z, Beare DM, Bindal N, Boutselakis H, Cole CG, Creatore C, Dawson E, Fish P, Harsha B, Hathaway C, Jupe SC, Kok CY, Noble K, Ponting L, Ramshaw CC, Rye CE, Speedy HE, Stefancsik R, Thompson SL, Wang S, Ward S, Campbell PJ, Forbes SA. COSMIC: the Catalogue Of Somatic Mutations In Cancer. Nucleic Acids Res. 2019 Jan 8;47(D1):D941–7.

46. Takeda AJ, Maher TJ, Zhang Y, Lanahan SM, Bucklin ML, Compton SR, Tyler PM, Comrie WA, Matsuda M, Olivier KN, Pittaluga S, McElwee JJ, Long Priel DA, Kuhns DB, Williams RL, Mustillo PJ, Wymann MP, Koneti Rao V, Lucas CL. Human PI3Kγ deficiency and its microbiota-dependent mouse model reveal immunodeficiency and tissue immunopathology. Nat Commun. Nature Publishing Group; 2019 Sep 25;10(1):4364–12.

47. Thian M, Hoeger B, Kamnev A, Poyer F, Köstel Bal S, Caldera M, Jiménez-Heredia R, Huemer J, Pickl WF, Groß M, Ehl S, Lucas CL, Menche J, Hutter C, Attarbaschi A, Dupré L, Boztug K. Germline biallelic PIK3CG mutations in a multifaceted immunodeficiency with immune dysregulation. Haematologica. Haematologica; 2020 Jan 30;:haematol.2019.231399.

48. Fruman DA, Rommel C. PI3K and cancer: lessons, challenges and opportunities. Nat Rev Drug Discov. 2014 Jan 31;13(2):140–56.

49. André F, Ciruelos E, Rubovszky G, Campone M, Loibl S, Rugo HS, Iwata H, Conte P, Mayer IA, Kaufman B, Yamashita T, Lu Y-S, Inoue K, Takahashi M, Pápai Z, Longin A-S, Mills D, Wilke C, Hirawat S, Juric D, SOLAR-1 Study Group. Alpelisib for PIK3CA-Mutated, Hormone Receptor-Positive Advanced Breast Cancer. N Engl J Med. Massachusetts Medical Society; 2019 May 16;380(20):1929–40.

50. Brown JR, Byrd JC, Coutre SE, Benson DM, Flinn IW, Wagner-Johnston ND, Spurgeon SE, Kahl BS, Bello C, Webb HK, Johnson DM, Peterman S, Li D, Jahn TM, Lannutti BJ, Ulrich RG, Yu AS, Miller LL, Furman RR. Idelalisib, an inhibitor of phosphatidylinositol 3 kinase p110δ, for relapsed/refractory chronic lymphocytic leukemia. Blood. 2014 Mar 10;123(22):3390–7.

51. Flinn IW, Kahl BS, Leonard JP, Furman RR, Brown JR, Byrd JC, Wagner-Johnston ND, Coutre SE, Benson DM, Peterman S, Cho Y, Webb HK, Johnson DM, Yu AS, Ulrich RG, Godfrey WR, Miller LL, Spurgeon SE. Idelalisib, a selective inhibitor of phosphatidylinositol 3-kinase-δ, as therapy for previously treated indolent non-Hodgkin lymphoma. Blood. 2014 Mar 10;123(22):3406–13.

52. Collier PN, Martinez-Botella G, Cornebise M, Cottrell KM, Doran JD, Griffith JP, Mahajan S, Maltais F, Moody CS, Huck EP, Wang T, Aronov AM. Structural basis for isoform selectivity in a class of benzothiazole inhibitors of phosphoinositide 3- kinase γ. J Med Chem. 2015 Jan 8;58(1):517–21.

53. Evans CA, Liu T, Lescarbeau A, Nair SJ, Grenier L, Pradeilles JA, Glenadel Q, Tibbitts T, Rowley AM, DiNitto JP, Brophy EE, O’Hearn EL, Ali JA, Winkler DG, Goldstein SI, O’Hearn P, Martin CM, Hoyt JG, Soglia JR, Cheung C, Pink MM, Proctor JL, Palombella VJ, Tremblay MR, Castro AC. Discovery of a Selective Phosphoinositide-3-Kinase (PI3K)-γ Inhibitor (IPI-549) as an Immuno-Oncology Clinical Candidate. ACS Med Chem Lett. American Chemical Society; 2016 Sep 8;7(9):862–7.

54. Safina BS, Elliott RL, Forrest AK, Heald RA, Murray JM, Nonomiya J, Pang J, Salphati L, Seward EM, Staben ST, Ultsch M, Wei B, Yang W, Sutherlin DP. Design of Selective Benzoxazepin PI3Kδ Inhibitors Through Control of Dihedral Angles. ACS Med Chem Lett. 2017 Sep 14;8(9):936–40.

55. Vadas O, Jenkins ML, Dornan GL, Burke JE. Using Hydrogen-Deuterium Exchange Mass Spectrometry to Examine Protein-Membrane Interactions. Meth Enzymol. Elsevier; 2017;583:143–72.

56. Dornan GL, Burke JE. Molecular Mechanisms of Human Disease Mediated by Oncogenic and Primary Immunodeficiency Mutations in Class IA Phosphoinositide 3-Kinases. Front Immunol. Frontiers; 2018;9:575.

57. Burke JE, Williams RL. Dynamic steps in receptor tyrosine kinase mediated activation of class IA phosphoinositide 3-kinases (PI3K) captured by H/D exchange (HDX-MS). Adv Biol Regul. 2013 Jan;53(1):97–110.

58. Burke JE, Perisic O, Masson GR, Vadas O, Williams RL. Oncogenic mutations mimic and enhance dynamic events in the natural activation of phosphoinositide 3-kinase p110α (PIK3CA). Proc Natl Acad Sci USA. 2012 Sep 18;109(38):15259– 64.

59. Burke JE, Vadas O, Berndt A, Finegan T, Perisic O, Williams RL. Dynamics of the phosphoinositide 3-kinase p110δ interaction with p85α and membranes reveals aspects of regulation distinct from p110α. 2011 Aug 10;19(8):1127–37.

60. Bruce I, Akhlaq M, Bloomfield GC, Budd E, Cox B, Cuenoud B, Finan P, Gedeck P, Hatto J, Hayler JF, Head D, Keller T, Kirman L, Leblanc C, Le Grand D, McCarthy C, O’Connor D, Owen C, Oza MS, Pilgrim G, Press NE, Sviridenko L, Whitehead L. Development of isoform selective PI3-kinase inhibitors as pharmacological tools for elucidating the PI3K pathway. Bioorganic & Medicinal Chemistry Letters. 2012 Sep 1;22(17):5445–50.

61. Knight Z, Gonzalez B, Feldman M, Zunder E, Goldenberg D, Williams O, Loewith R, Stokoe D, Balla A, Toth B, Balla T, Weiss W, Williams R, Shokat K. A pharmacological map of the PI3-K family defines a role for p110alpha in insulin signaling. Cell. 2006 Jun 19;125(4):733–47.

62. Knight SD, Adams ND, Burgess JL, Chaudhari AM, Darcy MG, Donatelli CA, Luengo JI, Newlander KA, Parrish CA, Ridgers LH, Sarpong MA, Schmidt SJ, Van Aller GS, Carson JD, Diamond MA, Elkins PA, Gardiner CM, Garver E, Gilbert SA, Gontarek RR, Jackson JR, Kershner KL, Luo L, Raha K, Sherk CS, Sung C-M, Sutton D, Tummino PJ, Wegrzyn RJ, Auger KR, Dhanak D. Discovery of GSK2126458, a Highly Potent Inhibitor of PI3K and the Mammalian Target of Rapamycin. ACS Med Chem Lett. American Chemical Society; 2010 Apr 8;1(1):39–43.

63. Venkatesan AM, Dehnhardt CM, Delos Santos E, Chen Z, Santos Dos O, Ayral-Kaloustian S, Khafizova G, Brooijmans N, Mallon R, Hollander I, Feldberg L, Lucas J, Yu K, Gibbons J, Abraham RT, Chaudhary I, Mansour TS. Bis(morpholino-1,3,5-triazine) derivatives: potent adenosine 5’-triphosphate competitive phosphatidylinositol-3-kinase/mammalian target of rapamycin inhibitors: discovery of compound 26 (PKI-587), a highly efficacious dual inhibitor. J Med Chem. 2010 Mar 25;53(6):2636–45.

64. Berndt A, Miller S, Williams O, Le DD, Houseman BT, Pacold JI, Gorrec F, Hon W-C, Liu Y, Rommel C, Gaillard P, Rückle T, Schwarz MK, Shokat KM, Shaw JP, Williams RL. The p110 delta structure: mechanisms for selectivity and potency of new PI(3)K inhibitors. Nature Chemical Biology. Nature Publishing Group; 2010 Feb;6(2):117–24.

65. Afonine PV, Moriarty NW, Mustyakimov M, Sobolev OV, Terwilliger TC, Turk D, Urzhumtsev A, Adams PD. FEM: feature-enhanced map. Acta Crystallogr D Biol Crystallogr. 2015 Mar;71(Pt 3):646–66.

66. Okkenhaug K. Signaling by the phosphoinositide 3-kinase family in immune cells. Annu Rev Immunol. 2013;31:675–704.

67. Sasaki T, Irie-Sasaki J, Horie Y, Bachmaier K, Fata JE, Li M, Suzuki A, Bouchard D, Ho A, Redston M, Gallinger S, Khokha R, Mak TW, Hawkins PT, Stephens L, Scherer SW, Tsao M, Penninger JM. Colorectal carcinomas in mice lacking the catalytic subunit of PI(3)Kgamma. Nature. Nature Publishing Group; 2000 Aug 24;406(6798):897–902.

68. Barbier M, Attoub S, Calvez R, Laffargue M, Jarry A, Mareel M, Altruda F, Gespach C, Wu D, Lu B, Hirsch E, Wymann MP. Tumour biology. Weakening link to colorectal cancer? Nature. Nature Publishing Group; 2001 Oct 25;413(6858):796–6.

69. Torres C, Mancinelli G, Cordoba-Chacon J, Viswakarma N, Castellanos K, Grimaldo S, Kumar S, Principe D, Dorman MJ, McKinney R, Hirsch E, Dawson D, Munshi HG, Rana A, Grippo PJ. p110γ deficiency protects against pancreatic carcinogenesis yet predisposes to diet-induced hepatotoxicity. Proc Natl Acad Sci USA. National Academy of Sciences; 2019 Jul 16;116(29):14724–33.

70. Dituri F, Mazzocca A, Lupo L, Edling CE, Azzariti A, Antonaci S, Falasca M, Giannelli G. PI3K class IB controls the cell cycle checkpoint promoting cell proliferation in hepatocellular carcinoma. Int J Cancer. John Wiley & Sons, Ltd; 2012 Jun 1;130(11):2505–13.

71. Edling CE, Selvaggi F, Buus R, Maffucci T, Di Sebastiano P, Friess H, Innocenti P, Kocher HM, Falasca M. Key role of phosphoinositide 3-kinase class IB in pancreatic cancer. Clin Cancer Res. 2010 Oct 15;16(20):4928–37.

72. Ge Y, He Z, Xiang Y, Wang D, Yang Y, Qiu J, Zhou Y. The identification of key genes in nasopharyngeal carcinoma by bioinformatics analysis of high-throughput data. Mol Biol Rep. 2019 Jun;46(3):2829–40.

73. Zhang P, Kang B, Xie G, Li S, Gu Y, Shen Y, Zhao X, Ma Y, Li F, Si J, Wang J, Chen J, Yang H, Xu X, Yang Y. Genomic sequencing and editing revealed the GRM8 signaling pathway as potential therapeutic targets of squamous cell lung cancer. Cancer Lett. 2019 Feb 1;442:53–67.

74. Nava Rodrigues D, Rescigno P, Liu D, Yuan W, Carreira S, Lambros MB, Seed G, Mateo J, Riisnaes R, Mullane S, Margolis C, Miao D, Miranda S, Dolling D, Clarke M, Bertan C, Crespo M, Boysen G, Ferreira A, Sharp A, Figueiredo I, Keliher D, Aldubayan S, Burke KP, Sumanasuriya S, Fontes MS, Bianchini D, Zafeiriou Z, Teixeira Mendes LS, Mouw K, Schweizer MT, Pritchard CC, Salipante S, Taplin M-E, Beltran H, Rubin MA, Cieslik M, Robinson D, Heath E, Schultz N, Armenia J, Abida W, Scher H, Lord C, D’Andrea A, Sawyers CL, Chinnaiyan AM, Alimonti A, Nelson PS, Drake CG, Van Allen EM, de Bono JS. Immunogenomic analyses associate immunological alterations with mismatch repair defects in prostate cancer. J Clin Invest. American Society for Clinical Investigation; 2018 Oct 1;128(10):4441–53.

75. Shu X, Gu J, Huang M, Tannir NM, Matin SF, Karam JA, Wood CG, Wu X, Ye Y. Germline genetic variants in somatically significantly mutated genes in tumors are associated with renal cell carcinoma risk and outcome. Carcinogenesis. 2018 May 28;39(6):752–7.

76. Wang J, Li M, Han X, Wang H, Wang X, Ma G, Xia T, Wang S. MiR-1976 knockdown promotes epithelial-mesenchymal transition and cancer stem cell properties inducing triple-negative breast cancer metastasis. Cell Death Dis. Nature Publishing Group; 2020 Jul 3;11(7):500–12.

77. Mandelker D, Gabelli SB, Schmidt-Kittler O, Zhu J, Cheong I, Huang C-H, Kinzler KW, Vogelstein B, Amzel LM. A frequent kinase domain mutation that changes the interaction between PI3Kalpha and the membrane. Proc Natl Acad Sci USA. National Acad Sciences; 2009 Oct 6;106(40):16996–7001.

78. Zhang X, Vadas O, Perisic O, Anderson KE, Clark J, Hawkins PT, Stephens LR, Williams RL. Structure of Lipid Kinase p110b/p85b Elucidatesan Unusual SH2- Domain-Mediated Inhibitory Mechanism. Mol Cell [Internet]. 2011 Apr 4;41(5):567–78. Available from: http://www.ncbi.nlm.nih.gov/entrez/query.fcgi?cmd=Retrieve&db=PubMed&dopt=Citation&list_uids=21362552

79. Perino A, Ghigo A, Ferrero E, Morello F, Santulli G, Baillie GS, Damilano F, Dunlop AJ, Pawson C, Walser R, Levi R, Altruda F, Silengo L, Langeberg LK, Neubauer G, Heymans S, Lembo G, Wymann MP, Wetzker R, Houslay MD, Iaccarino G, Scott JD, Hirsch E. Integrating Cardiac PIP(3) and cAMP Signaling through a PKA Anchoring Function of p110gamma. Mol Cell [Internet]. Elsevier Inc; 2011 Apr 8;42(1):84–95. Available from: http://www.ncbi.nlm.nih.gov/entrez/query.fcgi?cmd=Retrieve&db=PubMed&dopt=Citation&list_uids=21474070

80. Walser R, Burke JE, Gogvadze E, Bohnacker T, Zhang X, Hess D, Küenzi P, Leitges M, Hirsch E, Williams RL, Laffargue M, Wymann MP. PKCβ phosphorylates PI3Kγ to activate it and release it from GPCR control. Stephens L, editor. PLoS Biol. Public Library of Science; 2013;11(6):e1001587.

81. Berger I, Fitzgerald DJ, Richmond TJ. Baculovirus expression system for heterologous multiprotein complexes. Nat Biotechnol. Nature Publishing Group; 2004 Dec;22(12):1583–7.

82. Kozasa T. Purification of G protein subunits from Sf9 insect cells using hexahistidine-tagged alpha and beta gamma subunits. Methods Mol Biol. New Jersey: Humana Press; 2004;237:21–38.

83. Perez-Riverol Y, Csordas A, Bai J, Bernal-Llinares M, Hewapathirana S, Kundu DJ, Inuganti A, Griss J, Mayer G, Eisenacher M, Pérez E, Uszkoreit J, Pfeuffer J, Sachsenberg T, Yilmaz S, Tiwary S, Cox J, Audain E, Walzer M, Jarnuczak AF, Ternent T, Brazma A, Vizcaíno JA. The PRIDE database and related tools and resources in 2019: improving support for quantification data. Nucleic Acids Res. 2019 Jan 8;47(D1):D442–50.

84. Kabsch W. XDS. Acta Crystallogr D Biol Crystallogr. 2010 Feb;66(Pt 2):125–32.

85. McCoy AJ, Grosse-Kunstleve RW, Adams PD, Winn MD, Storoni LC, Read RJ. Phaser crystallographic software. J Appl Crystallogr. 2007 Jul 13;40(Pt 4):658–74.

86. Bohnacker T, Prota AE, Beaufils F, Burke JE, Melone A, Inglis AJ, Rageot D, Sele AM, Cmiljanovic V, Cmiljanovic N, Bargsten K, Aher A, Akhmanova A, Díaz JF, Fabbro D, Zvelebil M, Williams RL, Steinmetz MO, Wymann MP. Deconvolution of Buparlisib’s mechanism of action defines specific PI3K and tubulin inhibitors for therapeutic intervention. Nat Commun. Nature Publishing Group; 2017 Mar 9;8:14683.

87. Emsley P, Lohkamp B, Scott WG, Cowtan K. Features and development of Coot. Acta Crystallogr D Biol Crystallogr. 2010 Apr;66(Pt 4):486–501.

88. Afonine PV, Grosse-Kunstleve RW, Echols N, Headd JJ, Moriarty NW, Mustyakimov M, Terwilliger TC, Urzhumtsev A, Zwart PH, Adams PD. Towards automated crystallographic structure refinement with phenix.refine. Acta Crystallogr D Biol Crystallogr. 2012 Apr;68(Pt 4):352–67.

89. Chen VB, Arendall WB, Headd JJ, Keedy DA, Immormino RM, Kapral GJ, Murray LW, Richardson JS, Richardson DC. MolProbity: all-atom structure validation for macromolecular crystallography. Acta Crystallogr D Biol Crystallogr. 2010 Jan;66(Pt 1):12–21.

90. Sali A, Blundell TL. Comparative protein modelling by satisfaction of spatial restraints. Journal of Molecular Biology. 1993 Dec 5;234(3):779–815.

91. Sievers F, Wilm A, Dineen D, Gibson TJ, Karplus K, Li W, Lopez R, McWilliam H, Remmert M, Söding J, Thompson JD, Higgins DG. Fast, scalable generation of high-quality protein multiple sequence alignments using Clustal Omega. Mol Syst Biol. EMBO Press; 2011 Oct 11;7(1):539–9.

92. Case DA, Cheatham TE, Darden T, Gohlke H, Luo R, Merz KM, Onufriev A, Simmerling C, Wang B, Woods RJ. The Amber biomolecular simulation programs. J Comput Chem. 2005 Dec;26(16):1668–88.

93. Maier JA, Martinez C, Kasavajhala K, Wickstrom L, Hauser KE, Simmerling C. ff14SB: Improving the Accuracy of Protein Side Chain and Backbone Parameters from ff99SB. J Chem Theory Comput. 2015 Aug 11;11(8):3696–713.

94. Jorgensen WL, Chandrasekhar J, Madura JD, Impey RW, Klein ML. Comparison of simple potential functions for simulating liquid water. The Journal of Chemical Physics. American Institute of PhysicsAIP; 1998 Aug 31;79(2):926–35.

95. Miao Y, Feher VA, McCammon JA. Gaussian Accelerated Molecular Dynamics: Unconstrained Enhanced Sampling and Free Energy Calculation. J Chem Theory Comput. American Chemical Society; 2015 Aug 11;11(8):3584–95.

96. Pang YT, Miao Y, Wang Y, McCammon JA. Gaussian Accelerated Molecular Dynamics in NAMD. J Chem Theory Comput. American Chemical Society; 2017 Jan 10;13(1):9–19.

97. McGibbon RT, Beauchamp KA, Harrigan MP, Klein C, Swails JM, Hernández CX, Schwantes CR, Wang L-P, Lane TJ, Pande VS. MDTraj: A Modern Open Library for the Analysis of Molecular Dynamics Trajectories. Biophys J. 2015 Oct 20;109(8):1528–32.

## Supplemental references

1. Safina BS, Elliott RL, Forrest AK, Heald RA, Murray JM, Nonomiya J, Pang J, Salphati L, Seward EM, Staben ST, Ultsch M, Wei B, Yang W, Sutherlin DP. Design of Selective Benzoxazepin PI3Kδ Inhibitors Through Control of Dihedral Angles. ACS Med Chem Lett. 2017 Sep 14;8(9):936–40.

2. Furet P, Guagnano V, Fairhurst RA, Imbach-Weese P, Bruce I, Knapp M, Fritsch C, Blasco F, Blanz J, Aichholz R, Hamon J, Fabbro D, Caravatti G. Discovery of NVP-BYL719 a potent and selective phosphatidylinositol-3 kinase alpha inhibitor selected for clinical evaluation. Bioorganic & Medicinal Chemistry Letters. 2013 Jul 1;23(13):3741–8.

3. Zhang X, Vadas O, Perisic O, Anderson KE, Clark J, Hawkins PT, Stephens LR, Williams RL. Structure of Lipid Kinase p110b/p85b Elucidatesan Unusual SH2-Domain-Mediated Inhibitory Mechanism. Mol Cell [Internet]. 2011 Apr 4;41(5):567–78. Available from: http://www.ncbi.nlm.nih.gov/entrez/query.fcgi?cmd=Retrieve&db=PubMed&dopt=Citation&list_uids=21362552

4. Laskowski RA, Swindells MB. LigPlot+: multiple ligand-protein interaction diagrams for drug discovery. J Chem Inf Model. 2011 Oct 24;51(10):2778–86.

5. Heffron TP, Heald RA, Ndubaku C, Wei B, Augistin M, Do S, Edgar K, Eigenbrot C, Friedman L, Gancia E, Jackson PS, Jones G, Kolesnikov A, Lee LB, Lesnick JD, Lewis C, McLean N, Mörtl M, Nonomiya J, Pang J, Price S, Prior WW, Salphati L, Sideris S, Staben ST, Steinbacher S, Tsui V, Wallin J, Sampath D, Olivero AG. The Rational Design of Selective Benzoxazepin Inhibitors of the α-Isoform of Phosphoinositide 3-Kinase Culminating in the Identification of (S)-2-((2-(1-Isopropyl-1H-1,2,4-triazol-5-yl)-5,6- dihydrobenzo[f]imidazo[1,2-d][1,4]oxazepin-9-yl)oxy)propanamide (GDC-0326). J Med Chem. 2016 Feb 11;59(3):985–1002.

6. Somoza JR, Koditek D, Villaseñor AG, Novikov N, Wong MH, Liclican A, Xing W, Lagpacan L, Wang R, Schultz BE, Papalia GA, Samuel D, Lad L, McGrath ME. Structural, biochemical, and biophysical characterization of idelalisib binding to phosphoinositide 3-kinase δ. J Biol Chem. 2015 Mar 27;290(13):8439–46.

7. Afonine PV, Moriarty NW, Mustyakimov M, Sobolev OV, Terwilliger TC, Turk D, Urzhumtsev A, Adams PD. FEM: feature-enhanced map. Acta Crystallogr D Biol Crystallogr. 2015 Mar;71(Pt 3):646–66.

8. Knight Z, Gonzalez B, Feldman M, Zunder E, Goldenberg D, Williams O, Loewith R, Stokoe D, Balla A, Toth B, Balla T, Weiss W, Williams R, Shokat K. A pharmacological map of the PI3-K family defines a role for p110alpha in insulin signaling. Cell. 2006 Jun 19;125(4):733–47.

9. Knight SD, Adams ND, Burgess JL, Chaudhari AM, Darcy MG, Donatelli CA, Luengo JI, Newlander KA, Parrish CA, Ridgers LH, Sarpong MA, Schmidt SJ, Van Aller GS, Carson JD, Diamond MA, Elkins PA, Gardiner CM, Garver E, Gilbert SA, Gontarek RR, Jackson JR, Kershner KL, Luo L, Raha K, Sherk CS, Sung C-M, Sutton D, Tummino PJ, Wegrzyn RJ, Auger KR, Dhanak D. Discovery of GSK2126458, a Highly Potent Inhibitor of PI3K and the Mammalian Target of Rapamycin. ACS Med Chem Lett. American Chemical Society; 2010 Apr 8;1(1):39–43.

10. Masson GR, Burke JE, Ahn NG, Anand GS, Borchers C, Brier S, Bou-Assaf GM, Engen JR, Englander SW, Faber J, Garlish R, Griffin PR, Gross ML, Guttman M, Hamuro Y, Heck AJR, Houde D, Iacob RE, Jørgensen TJD, Kaltashov IA, Klinman JP, Konermann L, Man P, Mayne L, Pascal BD, Reichmann D, Skehel M, Snijder J, Strutzenberg TS, Underbakke ES, Wagner C, Wales TE, Walters BT, Weis DD, Wilson DJ, Wintrode PL, Zhang Z, Zheng J, Schriemer DC, Rand KD. Recommendations for performing, interpreting and reporting hydrogen deuterium exchange mass spectrometry (HDX-MS) experiments. Nat Methods. Nature Publishing Group; 2019 Jul;16(7):595–602.

